# Hydrothermal sulphur bacteria enhance mercury availability for coastal marine organisms

**DOI:** 10.1101/2022.04.26.489323

**Authors:** Eliot Ruiz, Thomas Lacoue-Labarthe, Maud Brault-Favrou, Pierre-Yves Pascal

## Abstract

The hydrothermal compounds massively released into the sea by the geothermal power plant of Bouillante (Guadeloupe, Lesser Antilles) support the growth of sulphur-oxidising bacteria like around black smokers. Opportunistic animals of the bay were previously shown to actively seek and consume the bacterial biofilm. To investigate the role of these bacterial mats in enhancing the transfer of Hg, a highly toxic heavy metal, its concentrations were assessed in sediments, bacteria, and 6 low trophic levels animals from the Bouillante Bay and a Control Site nearby containing only natural sources of Hg. Concentrations of Hg in all samples from Bouillante were greater than those from the Control Site (2 to 627 times higher). A comparison with the Hg concentrations reported in the literature for similar samples types reveals that they are abnormally high in most Bouillante samples. Although bacterial biofilm contained high levels of Hg, the link between bacteria contribution to diet and Hg concentration was more complex than expected, which might be due to interspecific differences in Hg uptake and elimination rates. Species not consuming bacteria (i.e. filter feeders) still presented higher levels of Hg, suggesting that significant amounts of Hg are released along with discharge waters from the Bouillante geothermal plant, and are integrated through diffusion. Differences in Hg concentrations between tissues do not indicate that Hg contained in samples from animals was under the form of MeHg, lowering the biomagnification potential of Hg in the Bouillante Bay trophic food web. Overall, the Bouillante case study emphasises for the first time an important role played by sulphur bacteria mats as a vector of hydrothermal inorganic Hg, and potentially other trace metals emitted in vents area, through dietary pathways.

## 1. Introduction

Due to the growing number of applications for the industrial development of metallic trace elements, their release into the environment is still increasing exponentially in some developing countries (Adeyanju & Okeke, 2019) and cause a great threat to human health and ecosystems. This major concern is reflected by the number of scientific papers dealing with them multiplicated by 35 in 20 years (Han et al., 2020). Among these elements, mercury (Hg) is considered the most bothering contaminant as it is highly toxic for all living organisms (Liu et al., 2012). Moreover, this metal could be transformed under its organic form methylmercury (MeHg), which is the most easily absorbed and retained form of Hg in animals. Consequently, MeHg raises major concerns since it is biomagnified in the food web, and can reach dangerous levels in top predators (Bosch et al., 2015).

Natural sources only account for a limited proportion (∼17%) of all Hg released in surface oceans compared to recent anthropogenic contributions since 1850 (∼70%; Amos et al., 2013). Globally, the main natural processes controlling Hg concentration in surface oceans are atmospheric deposition and oceanic evasion (Jiskra et al., 2021), but restricted sources such as river plumes with concentrated Hg or geothermal vents degassing inorganic Hg are locally significant (Amos et al., 2013). Indeed, they probably shaped the surrounding ecosystems at local scales by enhancing marine organisms’ exposure to Hg (e.g. Martins et al., 2001; Martins et al., 2006) and fostering specific local adaptation mechanisms with respect to Hg contamination (e.g. Liu et al., 2012; Vidal & Horne, 2003; Weis, 2002).

The Lesser Antilles constitute an active volcanic area with many active vent fields. Among these sites, the west coast of Basse-Terre in Guadeloupe presents several hydrothermal sources occurring at very shallow depths (Sanjuan & Bach, 1997), but also in streams pouring into the sea (Bagnato et al., 2009). This region is also characterized by high levels of natural Hg in sediments (Fabriol & Hazan, 1984; Morrison et al., 2015), and it was therefore hypothesised that shallow vent fields occurring there could be sources of Hg. In addition, these vents release great concentrations of sulphur compounds into the sea (Fabriol & Ouzounian, 1985), allowing the growth of many sulphur-bacteria species in the Bouillante Bay (P-Y. Pascal pers. obs.), potentially involved in the methylation of Hg. In addition, since 1986, the “Bouillante” geothermal plant constructed very close to the sea greatly enhanced the release of hydrothermal waters into the Bay. Indeed, this plant is unique: the part of hydrothermal waters that were not directly reinjected in wells is cooled with pumped seawater and directly rejected into the sea through a discharge channel. In comparison, other geothermal plants close to the sea either use cooling towers with recirculation or store wastewaters in lakes (Uihlein, 2018).

Therefore, the Bouillante Bay, as a very shallow hydrothermal system (depth lower than 25m), provide the unique opportunity to study many aspects linked to natural Hg, such as pathways of transfer to organisms, with limited resources due to their accessibility. Such very shallow vent fields are only found in 28 known locations worldwide, representing 4% of all known (i.e. active or inferred) active vent fields (Beaulieu & Szafranski, 2020). In addition, as prospection studies on Hg were carried out before its construction (Fabriol & Hazan, 1984), the Bouillante Bay represents a rare opportunity to study a potential recent increase in Hg concentration in the coastal environment. Moreover, the environmental conditions in the discharge channel support development of sulphur-oxidising bacteria whose mats growing on the rocky floor are eaten by opportunistic organisms (i.e. fishes and sea urchins) and locally increase their abundance (Pascal et al., 2017). This trophic function, known in natural hydrothermal vents (e.g. Cardigos et al., 2005), raises the question of the potential role of these sulphur-bacteria mats on the integration and transfer of Hg in the local trophic web of the Bouillante Bay.

To assess the impact of enhanced releases of hydrothermal waters in the Bouillante Bay on Hg accumulation and transfer in coastal organisms, total Hg (THg) concentrations were measured in sediments and organisms, including bacterial mats, sponges, bivalves, sea urchins and fishes, whose diet dependence to the bacteria and trophic levels differ among each other (Pascal et al., 2017). These data were compared with Hg concentrations in the same samples collected in a Control Site, far from the hydrothermal vents of Bouillante.

## 2. Materials and Methods

### 2.1. Geothermal plant

The study area is located on the west coast of Guadeloupe, which belongs to the active Lesser Antilles volcanic arc (Figure 1). The multiple hot springs and fumaroles characterising this region explain the name of the town there: Bouillante, which means “boiling” in French. Similarly to the well characterised Champagne Hot Springs of Dominica (Kleint et al., 2017), the bay of Bouillante and the surrounding coastline are known for their numerous very shallow springs producing numerous strings of gas bubbles and a clear hydrothermal fluid (Sanjuan & Brach, 1997).

**Figure 1.**
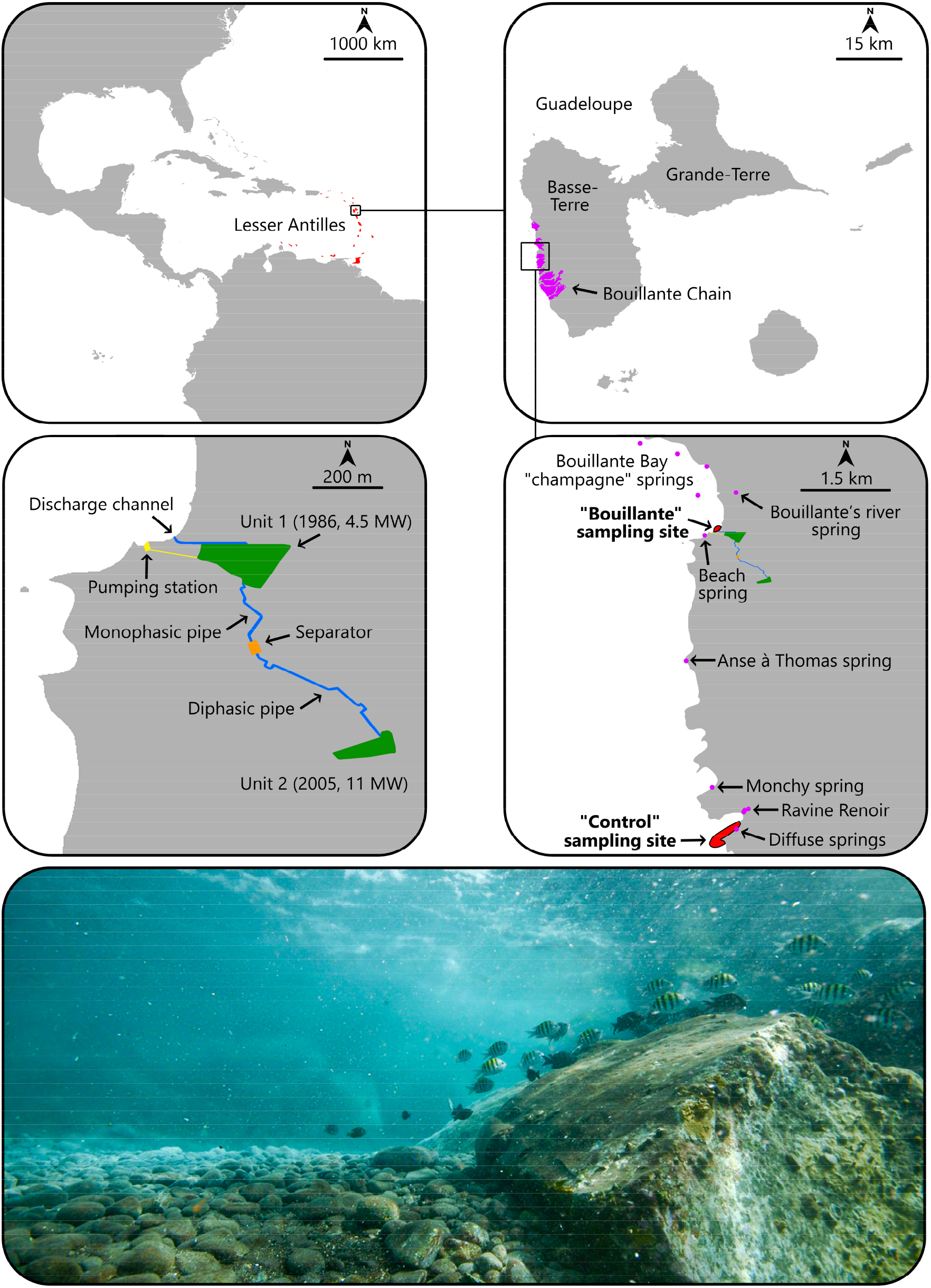
Map of the two sampling sites according to the location of the geothermal plant at different scales. The Bouillante Chain in purple on the upper right panel was represented according to Gadalia et al. (2019). The locations of all known springs pouring geothermal fluid in seawater represented in purple on the lower right panel were retrieved from Sanjuan & Bach (1997) and Bourdon et al. (2008) maps. A picture of the outlet of the discharge channel shows the bacterial white mat growing on the rocky pebbles in the background, and producing many clearly visible fragments ripped off by the strong current, which are eaten by *A. saxatilis* (left) and *A. bahianus* (centre).

This fluid originates from a 30,000,000 m3 reservoir (Sanjuan et al., 2004) formed by infiltrations of rainwater (40-45%) and seawater (55-60%; Sanjuan et al., 2000) in several fractured zones (tectonic activity) between 150m and 1100m (Lachassagne et al., 2010). After a period of reaction of more than a century with deep minerals (Sanjuan et al., 2004), a hot geothermal fluid (240-250°C; Sanjuan et al., 2000) emerges at the surface in both gaseous (20%) and liquid phases (80%) at an average temperature of 160-170°C (Dixit, 2014).

To exploit this green source of energy, one of the two France’s geothermal plants was put into service in 1986 in Bouillante, and now generates 6-7% (100-110 GWh per year) of the whole Guadeloupe Island electricity consumption (Demarcq et al., 2014). Multiple drillings were realised over the years to convey the high-pressure steam towards two interconnected turbines (Unit 1 & 2). The steam is partially condensed and partially released into the air, provoking a powerful smell of rotten eggs (due to H2S) near the units. The resulting water is then mixed with the liquid water separated from the vapour. To maintain the pressure in the aquifer, around 10% of the liquid water is reinjected while the rest is cooled down with pumped seawater (∼8380 m3.h-1), and then evacuated into the sea at a temperature of 40-45°C by a discharge channel (∼9000 m3.h-1; Géothermie Bouillante, 2018). The liquid flowing out in the sea through the channel consists of 22% of condensed vapour and 78% of separate liquid water (Géothermie Bouillante, 2018).

Among other chemicals, the geothermal fluid contains two sulphur molecules: sulphates in liquid water (SO4 = 13-17 mg.L-1) and hydrogen sulphide in vapour (H2S = 6 mg.L-1; Dixit, 2014). After mixing separate water and steam condensate, the SO4 (21 ± 2 mg.L-1; Dixit, 2014) and H2S (35-45 mg.L-1; Mas et al., 2006) content increase. To our knowledge, the chemical composition of the hydrothermal liquid mixed with seawater flowing in the discharge channel has not been measured yet.

### 2.2. Sampling sites and method

The Bouillante sampling site (GPS: 16.127964, -61.770353) is located near the shallow discharge channel (<1m) in which the current is strong (2.5 m^3^.s^−1^). At the mouth of the channel, there is a ∼45° pebble slope leading to a sedimentary bottom at a depth of 2m. The hot (40-45°C) discharge water flowing between the surface and the depth of the channel creates a strong thermocline. This limits the growth of the white mat covering the channel’s pebbles, even if fragments are continuously ripped off by the current and spread into the bay (**Figure 1**).

The thick mat (6-7mm) is made of benthic cyanobacteria, identified as *Plectonema sp*., covered with white filamentous bacteria of the genus *Thiomicrospira* (Pascal et al., 2017). These chemosynthetic bacteria use sulphur compounds (e.g. S^0^, H_2_S and S_2_O_3_^2-^) to create organic molecules in aerobic conditions thanks to their sulphur-oxidising activity (Takai et al., 2004). Therefore, they form the base of the food chain around deep hydrothermal vents, but can also be found in other marine ecosystems rich in sulphur compounds, since they are considered ubiquitous (Hansen & Perner, 2016).

The Control sampling site is situated on the southern side of the creek “Anse à la Barque” and around the “Pointe Diburque” (GPS: 16.087326, -61.768897), 4.5km away from Bouillante. This site contains many diffuse hot springs scattered in very shallow water, and also receives occasionally hydrothermal water from the multiple hot springs around the intermittent stream named “Ravine Renoir” nearby (Sanjuan & Bach, 1997; **Figure 1**).

Sampling took place on 07/14/17 and 07/18/17 and was carried out freediving. The white bacterial biofilm was collected through pebbles scraping. One sediment sample from Bouillante and three sediment samples from the Control site were collected in the first centimetres of the substrate at ∼2m depth. Sponges (*Aplysina fistularis* and *Iotrochota birotulata*), as well as sea urchins (*Diadema antillarum*) and bivalves (*Spondylus tenuis*) were collected by hand. Fishes (*Acanthurus bahianus* and *Abudefduf saxatilis*) were speared in the head, avoiding damaging the visceral mass and muscles. Ten specimens of each species were taken from each site.

### 2.3. Mercury concentration

The tissues chosen for our study were muscles in all animals except sponges, as well as gonads in urchins, since they are both edible and well-studied for Hg. The liver was also sampled in fishes since it allows to know if MeHg is the prevalent form of mercury (Scheuhammer et al., 2015). The whole branches were examined for sponges.

Fishes and sea urchins were measured (fishes: total length, urchins: test size) and weighed (wet weight). Ten samples of muscle were extracted for each species, as well as ten liver samples for each species of fish and ten gonad samples for sea urchins. Muscles extracted were protractor, retractor and compass elevator muscles for *D. antillarum*, adductor muscles for *S. tenuis* and fillets for fishes. Branches of sponges were cut into small pieces conserved as a whole. The 183 solid samples were then freeze-dried. Therefore, all concentrations of solid materials are based on dry weights.

Dry samples were reduced to a fine powder using a mortar and a pestle in order to homogenize them. THg concentrations (i.e. all mercury forms) were measured on a subsample of 0.05–50 mg dry powder (depending on the Hg concentrations) using an Advanced Mercury Analyser spectrophotometer (Altec AMA 254) able to detect 0.003ng to 150ng of Hg, as described in Chouvelon et al. (2009). Briefly, the samples were combusted under oxygen and the liberated Hg was analysed by atomic absorption spectrophotometry (Roos-Barraclough et al., 2002). For each sample, analyses were repeated two or three times, until the relative standard deviation between measures was <10%. Subsequently, the mean of the repeated Hg measurements was used for statistical analysis. To ensure the accuracy of measurements, a certified reference material, NRCC DOLT-3 (*Squalus acanthias* liver = 3.37 ± 0.14 μg.g^-1^), was analysed every 15 samples for recalibration of the analyser.

### 2.4. Data analyses

All statistical analyses were performed using R version 4.0.2. Since samples were not following a gaussian distribution, mean confidence intervals were calculated using 1 million iterations Bootstrap with the BCA (Biased-Corrected & Accelerated) method because it is considered the most conservative (Chou et al., 2006). The confidence intervals around mean ratios (i.e. Bouillante / Control) were computed with the same method, using a modified version of the function ci_mean_diff() from the *confintr* package to adapt it to ratios.

According to the validation or not of the conditions of use, means were compared using Student, Student with Welch correction or Wilcoxon tests. Differences between means with confidence interval and standardised effect size (i.e. Cohen’s d and Wilcoxon r) were calculated in case of significant p-values. Pairwise Wilcoxon tests were used to test for differences between Hg concentration in tissues for each species and among species. The Benjamini-Hochberg adjustment (1995) method was chosen to control the False Discovery Rate (FDR).

### 2.5. Risk assessment for Human consumers

A Maximum Safe Weekly Consumption (MSWC) was evaluated only for the tissues which are usually consumed by the local population. Thus, only *S. tenuis* muscle (Gobert & Reynal, 2002), *A. bahianus* muscle (Hawkins & Roberts, 2003) and *D. antillarum* gonads (Grisolia et al., 2012) were considered, since sponges, as well as livers, are not edible, and *A. saxatilis* is never used as food by local consumers due to its bad taste (Silvano & Begossi, 2012).

The MSWC was calculated on the basis of the Provisional Tolerable Weekly Intake (PTWI) given by the Joint Expert Committee on Food Additives (JECFA). The PTWI is 4 μg.kg^-1^ bw for inorganic mercury (Feeley et al., 2011) and 1.6 μg.kg^-1^ bw for methylmercury (Barlow et al., 2007). The average body weight (BW_ind_) in the Caribbean is 67.9kg (Walpole et al., 2012) and was used in the following formula:

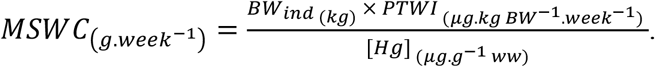

The Hg concentrations based on wet weight (ww) were converted to dry weight (dw) based concentrations using this formula:

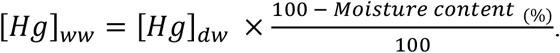

The conventional moisture content of 75% was used because it fitted well the values measured for close species, ranging from 73% to 77% (e.g. Elliott et al., 2015; Keshavarz & Jahromi, 2017; Szkoda et al., 2015).

## 3. Results

Among all 191 samples retrieved from the field, the mercury concentration could be successfully assessed for 189 of them. Indeed, the liver of the smallest *A. bahianus* (10 cm long) was too small to be analysed once dried, and the same problem occurred for the liver of one *A. saxatilis* specimen. Otherwise, the mean RSD for replicates in each sample was kept under 4%, which indicates a good accuracy of measurement.

### 3.1. Differences of Hg concentration among organisms

Regarding the Control Site, our analysis reveals intrinsic differences in Hg concentrations among the analysed organisms and tissues. Pairwise comparisons (Wilcoxon’s tests with FDR correction) between the concentrations measured in the samples from the Control Sites delineate 5 statistical groups, allowing to order sample types from the highest to the lowest total Hg concentrations: *A. bahianus* liver > *D. antillarum* gonad ∼ *A. fistularis* branch > *I. birotulata* branch > *A. saxatilis* liver ∼ *D. antillarum* muscle ∼ *S. tenuis* muscle ∼ *A. saxatilis* muscle > *A. bahianus* muscle (**Figure 2**).

**Figure 2.**
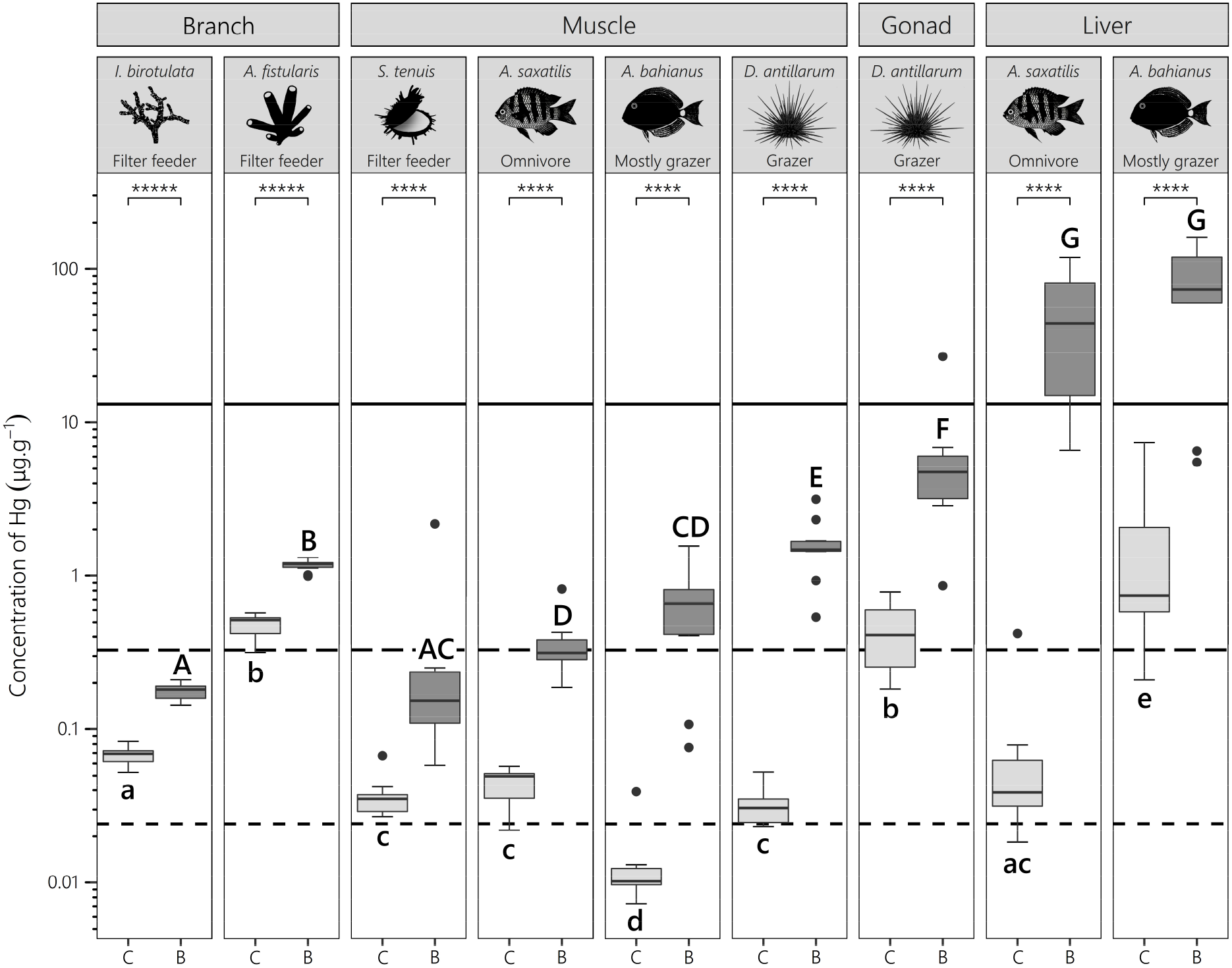
Hg concentrations (µg. g^-1^; dry weight) in marine organisms from the Control site (C, light grey) and Bouillante (B, dark grey) sites. The solid horizontal line represents the Hg concentration in the bacteria meadow ([Hg] = 13.01 µg.g^-1^). The dashed lines represent the Hg concentration in surface sediments (long-dash = Bouillante: [Hg] = 0.326 µg.g^-1^; short-dash = Control Site: [Hg] = 0.024 µg.g^-1^). The asterisks indicate a statistically significant difference (Student or Wilcoxon tests; p-value < 0.0001 = **** & p-value < 0.00001 = *****) between the two sampling sites. Letters regroup non-statistically different groups (Pairwise Wilcoxon’s Tests with FDR correction), the lower cases being used for the comparisons between Control samples and the upper cases for the comparisons between Bouillante samples. Concentrations were calculated based on dry weight.

The mean concentrations ranged from 0.013 μg.g^-1^ (95% CI [0.010, 0.024]) for *A. bahianus* muscle to 1.879 μg.g^-1^ (95% CI [0.866, 3.885]) for *A. bahianus* liver (i.e. 145-fold higher). The Hg concentrations measured in all samples but sponges, *A. bahianus* livers and *D. antillarum* livers, were similar or inferior to the sediment concentrations of the Control Site. This difference was maximal for *A. bahianus* liver, in which concentrations were 78 times higher than in the sediment, although there were large discrepancies between measurements for this sample type.

Regarding the Bouillante Bay, similarly to the results given by pairwise comparisons on the Control site’s Hg concentrations, sponges were not grouped together, as well as the gonads and muscle of *D. antillarum*. The opposed result was observed for the livers of fishes in which concentrations were non-statistically different, while they were highly divergent for the Control Site. Unlike the Control Site, Hg concentrations measured in Bouillante muscle samples were grouped in concomitant statistical groups (**Figure 2**).

The differences in Hg concentrations between samples were much higher in Bouillante than in the Control Site with 0.176 μg.g^-1^ (95% CI [0.163, 0.188]) in the branches of *I. birotulata* and 77.22 μg.g^-1^ (95% CI [45.70, 109.9]) in the liver of *A. bahianus* (i.e. 438-fold higher). The muscles of *S. tenuis, A. saxatilis* and *A. bahianus*, as well as the branches of *I. birotulata*, contained similar Hg concentrations to those found in the surface sediments of Bouillante. Mean Hg concentrations in fishes liver were 5.94 times higher than those measured in the bacterial mat for *A. bahianus*, and 4.10 times higher for *A. saxatilis*. The concentrations for the other sample types were situated between the concentrations found in the sediments and those found in the bacterial mat (**Table S1**).

### 3.2. Differences of Hg concentrations between sites

Hg concentrations were drastically higher in Bouillante than in the Control Site, whatever the sample type. All statistical tests (i.e. Student or Wilcoxon tests) were significant and standardised effect sizes (i.e. Cohen’s D or Wilcoxon’s R) indicated a large magnitude of effect (**Table S2**). Indeed, the predicted differences between both sites (Bouillante – Control) by Student or Wilcoxon tests were always positive, ranging from + 0.11 μg.g^-1^ (95% CI [0.09, 0.13]) for *Iotrochota birotulata* branches, to + 70.9 μg.g^-1^ (95% CI [52.6, 118]) for *Acanthurus bahianus* liver (**Table S2**).

This change was expressed as the ratio between both concentrations (**Figure 3**) allowing to assess the magnitude of the contamination increase. The Hg concentration in surficial sediment at 2m depth was much higher close to the discharge channel, with concentrations 13.63-fold higher in Bouillante than in the Control site. Interestingly, both *I. birotulata* and *A. fistularis* displayed the lowest Hg ratio with sponge branch only 2.64-fold (95% CI [2.37, 2.96]) and 2.44-fold (95% CI [2.20, 2.81]) more contaminated in Bouillante than in Control Site, respectively (**Table S1**). Considering the Hg ratio in muscle samples, *A bahianus* (× 48.51, 95% CI [24.96, 78.77]) and *D. antillarum* (× 49.75, 95% CI [36.50, 68.34]) displayed the highest increase of contamination in Bouillante, compared to *S. tenuis* (× 9.71, 95% CI [3.69, 30.02]) and *A. saxatilis* (× 8.38, 95% CI [6.43, 12.78]). Contrasting to this, regarding fish liver, *A. saxatilis* (× 626.82, 95% CI [191.24, 1589.50]) showed the highest Hg contamination raise compared to *A. bahianus* (× 41.09, 95% CI [16.22, 99.77]), suggesting that both tissues reflect contrasting bioaccumulation pathways occurring in Bouillante.

**Figure 3.**
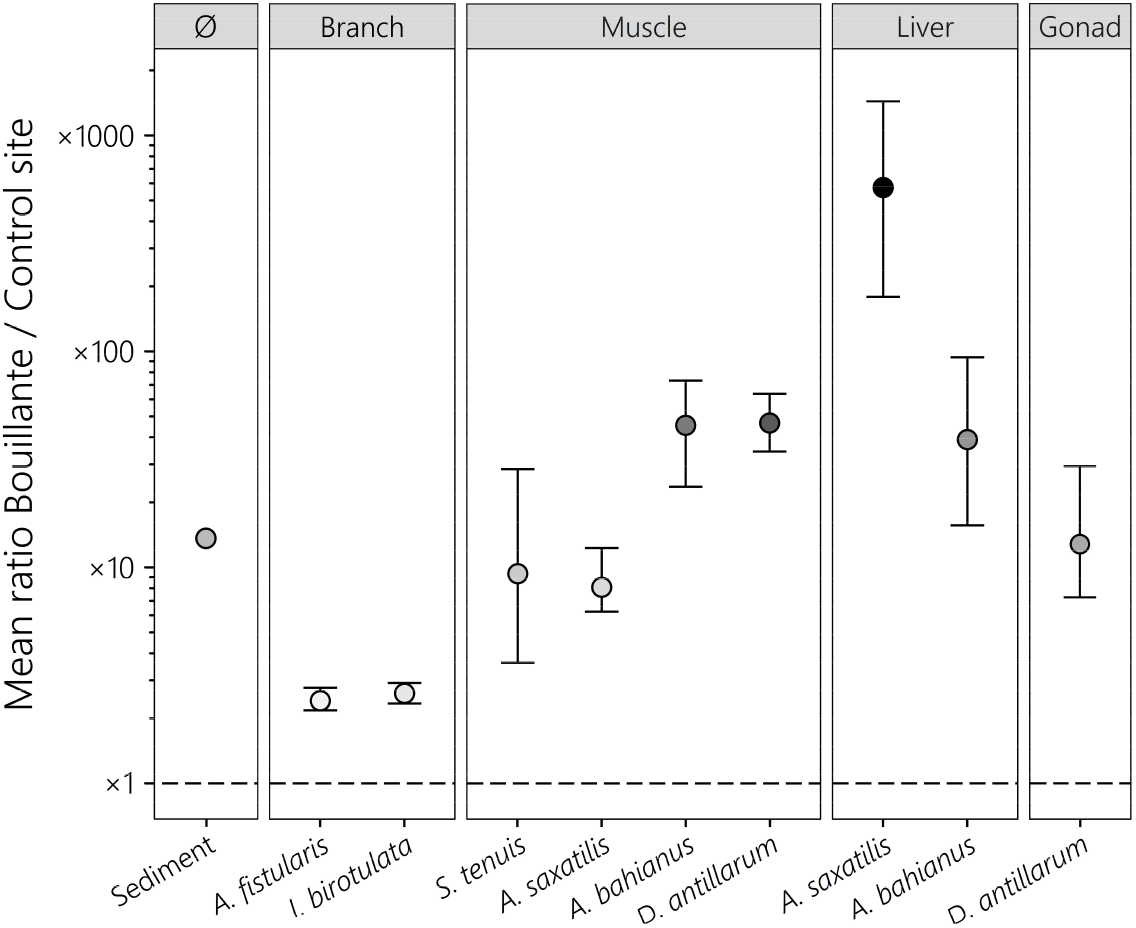
Mean ratio between the Hg concentration of Bouillante and the Control Site, enabling to evaluate the effect of the geothermal plant independently of initial differences between types, species and tissues. The vertical bars represent the 95% bootstrapped confidence interval around the mean ratio (iterations = 1,000,000). Ratios were established using concentrations in µg.g^-1^ (dry weight). The different shades of grey are proportional to the mean ratio. The dashed line corresponds to equal concentrations in Bouillante and the Control Site. Concentrations were calculated based on dry weight.

### 3.4. Risk assessment for human consumers

While the Hg speciation in the respective tissues was not assessed in this study, the MSWC values were calculated considering both Provisional Tolerable Weekly Intake (PTWI) for inorganic Hg (Hg_inorg_) and MeHg. The MSWC with respect to seafood items from the Control Site ranged from 2.43 kg.week^-1^ (95% CI [1.85, 3.35] for *D. antillarum* gonads to 83.6 kg.week^-1^ (95% CI [45.3, 108.7]) for *A. bahianus* muscle, with the assumption that accumulated Hg was under inorganic form. The concentrations of Hg in Bouillante organisms being higher than in the Control Site, the MSWC was therefore lower for all samples ranging from 167 g.week^-1^ (95% CI [77.9, 282.9]) of *D. antillarum* gonad to 3.1 kg.week^-1^ (95% CI [0.95, 7.8]) of *S. tenuis* muscle.

The same tendencies appeared in the MeHg scenario, although all MSWC were approximately divided by 2.5 (**Table S3**).

## 4. Discussion

### 4.1. Mercury in sediments

Prospection studies preceding the construction of the Bouillante geothermal plant in 1986 interestingly revealed that THg in shallow surface sediments near the future discharge channel (0.065 μg.g^-1^) was already higher in Bouillante than in the Control Site (0.040 μg.g^-1^; Fabriol & Hazan, 1984). This could be due to the presence of a higher number of scattered shallow bubbling springs in the sediment bed near the Bouillante sampling site than in the Control sampling site (P-Y. Pascal pers. obs.).

In addition, the concentrations measured in shallow surface sediments on the Control Site (i.e. 0.024 µg.g^-1^, 95% CI [45.70, 109.9]) were ∼1.7 times lower than the values reported in 1984 (Fabriol & Hazan, 1984). Contrasting to this, Hg concentrations in sediment from Bouillante have become 5 times higher over the last 4 decades. Present values for the Control Site are also similar to those reported in other probable Hg-poor sites (i.e. 0.001-0.041 μg.g^-1^; Morrison et al., 2015; **Table 1**), but concentrations in Bouillante largely exceed these values (0.326 µg.g^-1^). This result could be explained by the enhanced release of Hg-rich hydrothermal waters through the discharge channel (∼620 m^3^.h^-1^; Géothermie Bouillante, 2018). Indeed, hydrothermal waters from the Bouillante chain’s hot springs are characterised by abnormally high concentrations of Hg compared to other active volcanic areas (i.e. 10 – 376 ng.L^-1^; Bagnato et al., 2009). However, since low-density brackish wastewaters form a slick at the surface of the sea visible 900m away from the discharge channel (Caraïbes Environnement, 2011), with low dilution in a 300m radius around the source (PARETO – IMPACTMER, 2009) and a strong thermocline, it is unlikely that significant amount of dissolved Hg reach the sediment bed where we collected sediment samples in Bouillante. Therefore, it is also improbable that large amounts of sulphur-rich discharge water could diffuse into sediments and support sulphate-reducing bacteria, the main producers of MeHg (Parks et al., 2013). However, bacterial mat fragments and particulate Hg forming a sort of “marine snow” rapidly settling on the substrate near the mouth of the channel could be sequestered in sediments and increase its Hg content.

**Table 1.**
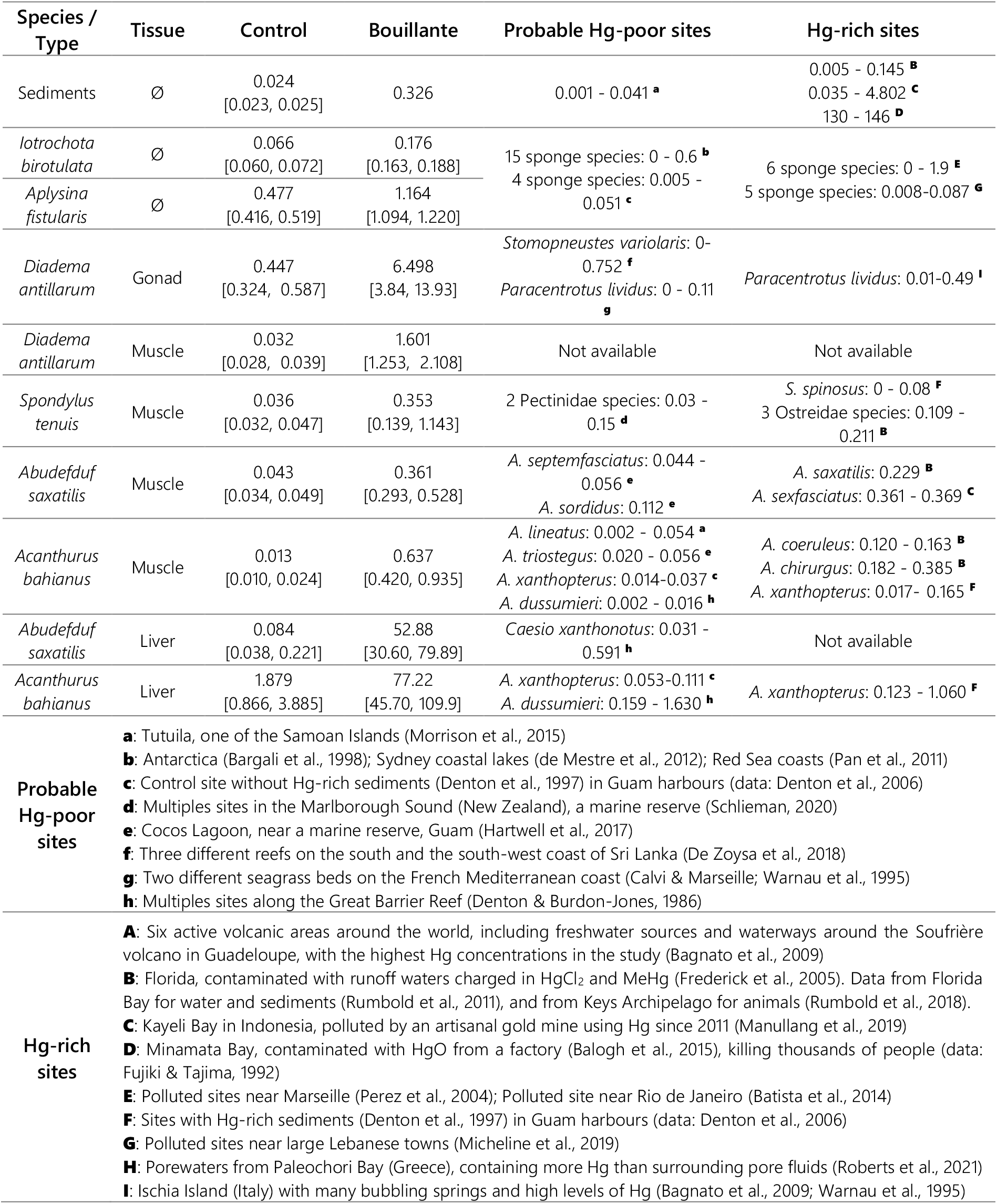
Comparison between Hg concentrations measured in this study and Hg concentrations measured in other sites worldwide differing in Hg contamination levels. Only species with close biology, ecology and trophic levels were compared. The range (min – max) of Hg concentrations between sample types in other articles is reported in the last two columns instead of the mean and their 95% robust CI in the first two columns, since not enough data was available for calculations of these statistics in most cases. All concentrations are based on dry weight and reported μg.g^-1^. Concentrations given in wet weight in other articles were converted in dry-weight based concentrations using a conventional 75% moisture content if not given.

| Species / Type | Tissue | Control | Bouillante | Probable Hg-poor sites | Hg-rich sites |
| --- | --- | --- | --- | --- | --- |
| Sediments | Ø | 0.024<br>[0.023, 0.025] | 0.326 | 0.001 - 0.041 <sup>a</sup> | 0.005 - 0.145 <sup>B</sup><br>0.035 - 4.802 <sup>C</sup><br>130 - 146 <sup>D</sup> |
| *Iotrochota birotulata* | Ø | 0.066<br>[0.060, 0.072] | 0.176<br>[0.163, 0.188] | 15 sponge species: 0 - 0.6 <sup>b</sup><br>4 sponge species: 0.005 - 0.051 <sup>c</sup> | 6 sponge species: 0 - 1.9 <sup>E</sup><br>5 sponge species: 0.008-0.087 <sup>G</sup> |
| *Aplysina fistularis* | Ø | 0.477<br>[0.416, 0.519] | 1.164<br>[1.094, 1.220] |  |  |
| *Diadema antillarum* | Gonad | 0.447<br>[0.324, 0.587] | 6.498<br>[3.84, 13.93] | *Stomopneustes variolaris*: 0-0.752 <sup>f</sup><br>*Paracentrotus lividus*: 0 - 0.11 <sup>g</sup> | *Paracentrotus lividus*: 0.01-0.49 <sup>I</sup> |
| *Diadema antillarum* | Muscle | 0.032<br>[0.028, 0.039] | 1.601<br>[1.253, 2.108] | Not available | Not available |
| *Spondylus tenuis* | Muscle | 0.036<br>[0.032, 0.047] | 0.353<br>[0.139, 1.143] | 2 Pectinidae species: 0.03 - 0.15 <sup>d</sup> | *S. spinosus*: 0 - 0.08 <sup>F</sup><br>3 Ostreidae species: 0.109 - 0.211 <sup>B</sup> |
| *Abudefduf saxatilis* | Muscle | 0.043<br>[0.034, 0.049] | 0.361<br>[0.293, 0.528] | *A. septemfasciatus*: 0.044 - 0.056 <sup>e</sup><br>*A. sordidus*: 0.112 <sup>e</sup> | *A. saxatilis*: 0.229 <sup>B</sup><br>*A. sexfasciatus*: 0.361 - 0.369 <sup>C</sup> |
| *Acanthurus bahianus* | Muscle | 0.013<br>[0.010, 0.024] | 0.637<br>[0.420, 0.935] | *A. lineatus*: 0.002 - 0.054 <sup>a</sup><br>*A. triostegus*: 0.020 - 0.056 <sup>e</sup><br>*A. xanthopterus*: 0.014-0.037 <sup>c</sup><br>*A. dussumieri*: 0.002 - 0.016 <sup>h</sup> | *A. coeruleus*: 0.120 - 0.163 <sup>B</sup><br>*A. chirurgus*: 0.182 - 0.385 <sup>B</sup><br>*A. xanthopterus*: 0.017- 0.165 <sup>F</sup> |
| *Abudefduf saxatilis* | Liver | 0.084<br>[0.038, 0.221] | 52.88<br>[30.60, 79.89] | *Caesio xanthonotus*: 0.031 - 0.591 <sup>h</sup> | Not available |
| *Acanthurus bahianus* | Liver | 1.879<br>[0.866, 3.885] | 77.22<br>[45.70, 109.9] | *A. xanthopterus*: 0.053-0.111 <sup>c</sup><br>*A. dussumieri*: 0.159 - 1.630 <sup>h</sup> | *A. xanthopterus*: 0.123 - 1.060 <sup>F</sup> |
| Probable Hg-poor sites |  | <sup>a</sup>: Tutuila, one of the Samoan Islands (Morrison et al., 2015)<br><sup>b</sup>: Antarctica (Bargali et al., 1998); Sydney coastal lakes (de Mestre et al., 2012); Red Sea coasts (Pan et al., 2011)<br><sup>c</sup>: Control site without Hg-rich sediments (Denton et al., 1997) in Guam harbours (data: Denton et al., 2006)<br><sup>d</sup>: Multiples sites in the Marlborough Sound (New Zealand), a marine reserve (Schlieman, 2020)<br><sup>e</sup>: Cocos Lagoon, near a marine reserve, Guam (Hartwell et al., 2017)<br><sup>f</sup>: Three different reefs on the south and the south-west coast of Sri Lanka (De Zoysa et al., 2018)<br><sup>g</sup>: Two different seagrass beds on the French Mediterranean coast (Calvi & Marseille; Warnau et al., 1995)<br><sup>h</sup>: Multiples sites along the Great Barrier Reef (Denton & Burdon-Jones, 1986) |  |  |  |
| Hg-rich sites |  | <sup>A</sup>: Six active volcanic areas around the world, including freshwater sources and waterways around the Soufrière volcano in Guadeloupe, with the highest Hg concentrations in the study (Bagnato et al., 2009)<br><sup>B</sup>: Florida, contaminated with runoff waters charged in HgCl<sub>2</sub> and MeHg (Frederick et al., 2005). Data from Florida Bay for water and sediments (Rumbold et al., 2011), and from Keys Archipelago for animals (Rumbold et al., 2018).<br><sup>C</sup>: Kayeli Bay in Indonesia, polluted by an artisanal gold mine using Hg since 2011 (Manullang et al., 2019)<br><sup>D</sup>: Minamata Bay, contaminated with HgO from a factory (Balogh et al., 2015), killing thousands of people (data: Fujiki & Tajima, 1992)<br><sup>E</sup>: Polluted sites near Marseille (Perez et al., 2004); Polluted site near Rio de Janeiro (Batista et al., 2014)<br><sup>F</sup>: Sites with Hg-rich sediments (Denton et al., 1997) in Guam harbours (data: Denton et al., 2006)<br><sup>G</sup>: Polluted sites near large Lebanese towns (Micheline et al., 2019)<br><sup>H</sup>: Porewaters from Paleochori Bay (Greece), containing more Hg than surrounding pore fluids (Roberts et al., 2021)<br><sup>I</sup>: Ischia Island (Italy) with many bubbling springs and high levels of Hg (Bagnato et al., 2009; Warnau et al., 1995) |  |  |  |

### 4.2. Mercury in bacteria

The white bacterial mat is exclusively covering the rocky pebbles of the shallow discharge channel and is made of filamentous *Plectonema sp*. cyanobacteria (Oscillatoriaceae) associated with the sulphur-oxidising *Thiomicrospira sp*. gammaproteobacteria (Pascal et al., 2017). Associations between Oscillatoriaceae and sulphur-oxidising bacteria were already reported in mats over sulphur-rich sediments (Stolz, 1985; Mir et al., 1991; Guézennec et al., 2011) since bacteria benefits from the O_2_ released by the cyanobacteria. The development of *Thiomicrospira* in the discharge channel is clearly linked with the Bouillante plant activity, since this obligate sulphur-oxidising bacteria (Takai et al., 2004) quickly disappears when discharge waters stop flowing during maintenance work, cutting off the abundant sulphur supply (Pascal et al., 2017).

The high concentration of Hg measured in the bacterial mats (13.01 µg.g^-1^) growing on the rocky pebbles (**Figure 2**) is likely caused by the probable release of Hg-rich discharge waters at high rates, since the upward migration of Hg^0^ would not reach the mats growing on rocky pebbles only. The dissolved inorganic Hg can easily diffuse through the thin lipid bilayer (Hsu-Kim et al., 2013), and particulate HgS (from condensed steam) can also enter the cell thanks to its strong affinity with thiol groups (Schaefer & Morel, 2009). Therefore, *Thiomicrospira* secretes their own exocellular polysaccharide (EPS) to compensate for the lack of the detoxifying *mer* system (Cruz, 2014), and also benefits from large EPS quantities excreted by *Plectonema* (Guézennec et al., 2011). Negatively charged groups in polysaccharides bind to Hg^2+^ (Volesky, 1990), and neutral Hg molecules are blocked by electrostatic interactions (e.g. >80% HgCl_2_ blocked by *Thiomicrospira* EPS; Cruz, 2014; Tan et al., 2016), explaining highly elevated concentrations in Bouillante’s bacterial mats.

As none *Thiomicrospira* strains (NCBI, 2022), and generally no Gammaproteobacteria (Podar et al., 2015; Gionfriddo et al., 2016), possess the hgcAB gene pair coding for proteins responsible of Hg methylation (Parks et al., 2013), it is unlikely that bacterial mats are a source of MeHg in Bouillante.

### 4.3. Mercury in animals of the Control Site

Concentrations of Hg in the Control Site seem consistent with previous values reported for the same species, or close species in terms of biology, ecology and trophic level, in other study zones with probably low levels of Hg (**Table 1**). They were also similar to those reported in Hg-enriched sites in some cases: for example, gonads of *D. antillarum* contain similar concentrations to those of *Paracentrotus lividus* in Ischia Island (Italy), indicating similarities between both sites (Warnau et al., 1995; Bagnato et al., 2009). Large differences in Hg concentration between samples types of the Control Site were measured, with significant differences between species as well as between tissues within species, indicating basal differences in Hg accumulation (**Figure 2**).

Regarding sponges, Hg concentrations recorded in *I. birotulata* and *A. fistularis* were in the range of values reported in sponges from non-polluted areas (**Table 1**). Interestingly, Hg concentrations in *A. fistularis* were on average 7 times more elevated than in *I. birotulata*. This difference could be linked to their different shapes (Lacoue-Labarthe et al., 2016), their differences of microbial abundance (Pawlik et al., 2015) or their differences in microbial communities (e.g. symbiotic *Synechococcus* of *A. fistularis* can act on HgCl_2_; Erwin & Thacker, 2007; Lefebvre et al., 2007). In addition, the elevated filtration rate of *A. fistularis* (Weisz, et al., 2008) is probably inferior in *I. birotulata* due to the numerous small polyps (Zoanthidae: *Parazoanthus swiftii*; Crocker & Reiswig, 1981) often settling in its tissue and reducing the water flow through the sponge (Lewis, 1982).

In fish, both liver and muscle exhibit specific concentrations and Hg speciation, as a consequence of tissue components, detoxification mechanisms, but also depending on bioaccumulation pathways (waterborne *vs*. diet), food regime and proportion of Hg species in food items (Maury-Brachet et al., 2006; Wang & Wong, 2003). Interestingly, *A. bahianus* showed the lowest and the highest Hg concentrations in the muscle and the liver, respectively, likely because its benthivorous/periphytophagous diet favours the ingestion of inorganic forms of Hg (48% to 72% of Hg in their food) that preferentially accumulates in the liver and kidneys (Maury-Brachet et al., 2006). On the contrary, the planktivorous *A. saxatilis*, which has a higher trophic level (δ^15^N = 5.70 ± 0.31‰) than *A. bahianus* (δ^15^N = 4.89 ± 0.10‰; Pascal et al., 2017), likely contains more MeHg, since concentrations in its muscle and liver are similar (Scheuhammer et al., 2015). The comparison with other probable Hg-poor sites reveals similar tendencies for fishes’ liver and muscle (**Table 1**) indicating a classical accumulation of Hg in the tissue of these fishes, with differences between species probably explained by their different diet.

### 4.4. Mercury in animals of Bouillante

Regarding animals sampled in the Bouillante Bay, our study reveals a large and significant increase of the Hg content in all samples (**Figure 2 & Table S2**), whose magnitude depends on the species as well as on the tissue considered within each species (**Figure 3**). Clearly, these Hg concentrations demonstrate a Hg contamination on this site that can be linked with the release of discharge hydrothermal water by the Bouillante power plant, as shown by the increase of Hg content in sediments from 1984 to our study. In addition, we assume that the studied species reflected the local contamination as their ranges of movements are considered to be lower than 400m (Chapman & Kramer, 2000; Pascal et al., 2017). Moreover, this contamination appears important as Hg concentrations in Bouillante samples were generally much higher than those from close species in probable Hg-poor sites and even Hg-rich sites (**Table 1**). For example, Hg in livers of *A. bahianus* is ∼80-fold higher than in livers of *Acanthurus xanthopterus* from Hg-enriched harbours in Guam (Denton et al., 1997; Denton et al., 2006).

Interestingly, previous isotopic signature analyses (i.e. δ^15^N, δ^13^C & δ^34^S) showed that fish (*A. bahianus & A. saxatilis*) and sea urchin (*D. antillarum*), contrasting to filter feeders (i.e. *I. birotulata, A. fistularis & S. tenuis*), actively seek (i.e. greater abundance) and consume the bacterial mats growing in the channel (Pascal et al., 2017), suggesting a potential role of bacteria in the integration and transfer of Hg only through these opportunistic species. Our results indicate that bacterial consumption is indeed a significant pathway of Hg bioaccumulation in *A. bahianus, A. saxatilis* and *D. antillaru*m, since the ratios of increase in their tissues were elevated (**Figure 3**). However, filter feeders not known to consume bacterial mats still contain ∼2.5 (sponges) to ∼10 (*S. tenuis*) times more Hg in Bouillante than in the Control Site. For example, *A. saxatilis* has a slightly lower mean ratio than *S. tenuis* (**Figure 5**), while the latter is not known to consume bacterial mats contrarily to *A. saxatilis* (∼27% of the diet in Bouillante; Pascal et al., 2017), possibly because of active removing of bacterial mats fragments from the gills as shown for mussels (Trager & DeNiro, 1990). Since, the dissolved uptake rate of inorganic mercury (Hg^2+^) is generally much higher in bivalves (3.5 – 32.8 L.g^-1^.d^-1^) than in fish (0.038 – 0.195 L.g^-1^.d^-1^; Wang & Wong, 2003; Pickhardt et al., 2006; Pan & Wang, 2011), *S. tenuis* would consequently gain a higher amount of Hg by direct diffusion through its gills than *A. saxatilis*. Generally, elevated concentrations in filter feeders support the hypothesis that discharge waters contain higher concentrations of Hg than in the Control Site, indicating a second potential pathway of Hg integration into the Bouillante Bay animals through diffusion.

Moreover, the liver/muscle ratio from fishes of Bouillante much lower than **⅓** clearly indicates that Hg is not under a methylated form in these organisms (Scheuhammer et al., 2015), suggesting that the dominant Hg form in the bacterial mat is not MeHg. Unlike the Control Site, the liver/muscle ratio of both species of fishes is highly elevated in Bouillante (**Figure 1**), since that of *A. saxatilis* (C = 1.9 & B = 146.48) becomes similar to that of *A. bahianus* (C = 144.5 & B = 121.2), other *Acanthurus* (**Table 1**), and in general benthivorous/periphytophagous fishes (Cidziel et al., 2002; Régine et al., 2006). Indeed, while *A. bahianus* is expected to consume more bacterial mats, the mean ratio of increase in *A. saxatilis* liver is 15 times higher, and the absolute concentrations in livers are not statistically different between each species. This result confirms the partial diet shift showed by Pascal et al. (2017), from planktonic preys naturally containing mostly MeHg (i.e. accumulated in muscles), to bacterial mats containing mostly inorganic Hg (i.e. stored in liver and kidneys; Régine et al. 2006). Such rapid increase of Hg in the livers of *A. saxatilis* from Bouillante might be explained by a detoxifying system for inorganic Hg less efficient than *A. bahianus* which is naturally more exposed to inorganic Hg from food. For example, it has been shown that multiple Hg-binding proteins (e.g. metallothionein), which is one of the detoxification strategies along with Hg elimination in fish (Pan & Wang, 2012), are less efficient in carnivorous than in herbivorous fish species (Vieira et al., 2017).

Finally, as for *S. tenuis vs. A. saxatilis* and *A. bahianus vs. A. saxatilis*, Hg concentrations in sea urchins’ muscles also demonstrate that Hg concentration in muscle does not seem to proportionally increase with bacteria contribution to diet as initially expected (**Figure 5**). Indeed, *D. antillarum*, which generally consumes greater amounts of bacterial mats than *A. bahianus* (Pascal et al., 2017), has a lower ratio of increase in Hg (**Figure 5**), even if absolute concentrations in its muscle are significantly greater (**Figure 1**). This could be the result of much faster elimination rates of *D. antillarum* over *A. bahianus*, which has already been observed for other echinoderm and fish species (Bjerregaard & Møller, 2021). Concerning *D. antillarum* gonads, their significantly higher Hg concentration than in muscles in both sites reveals that Hg is preferentially accumulated in gonads (**Figure 1**). This is a new result since Hg concentration in muscle was not previously measured in sea urchins to our knowledge. Unlike fishes, the mean ratio of increase was 3.4 times higher in *D. antillarum* Aristotle’s lantern muscles (i.e. masticatory apparatus) than in its gonads, which might be linked to the great importance of diet for Hg integration in this species.

### 4.5. Ecological and human consequences

MSWC in the three edible tissues surveyed in this study do not present a large risk for humans in both probable Hg_inorg_ and improbable MeHg scenarios (**Figure 4**). For example, when considering Hg as inorganic, consuming ∼34 *A. bahianus* (1.7 kg.week^-1^, 95% CI [1.2, 2.6]) and ∼186 *S. tenuis* (3.1 kg.week^-1^, 95% CI [1.0, 7.8]) from Bouillante per week would be necessary to ingest harmful doses of Hg (**Table S3**). However, while *D. antillarum* is rarely consumed in Guadeloupe due to its very painful sting (Coppard & Campbell, 2004), the maximal dose of ∼21 adult urchins’ gonads per week (**Table S3**) when considering Hg as inorganic could well be crossed for the highly prized and similarly sized white urchin (*Tripneustes ventricosus*) which also lives in the Bouillante Bay (Colombet, 2021). Moreover, direct predators of *A. bahianus* and *A. saxatilis* such as *Sphyraena barracuda, Caranx sp*. or *Lutjanus sp*. (Miller et al., 1971) which are consumed in Guadeloupe and present near the discharge channel (P-Y. Pascal pers. obs.) could accumulate harmful levels of Hg. Besides, the MSWC of these predators’ muscles are low in the Florida Keys, notably contaminated with inorganic Hg (Frederick et al., 2005), especially for *Sphyraena barracuda* (minimum MSWC = 80 g.week^-1^; Rumbold et al., 2018). Generally, both diffusive and dietary Hg integration pathways probably occurring near the mouth of the discharge channel could have contaminated all low trophic levels organisms living nearby, and this Hg could then be biomagnified in the Bouillante Bay (e.g. Lacoue-Labarthe et al., 2016). Even burrowing organisms such as nematodes that do not consume bacterial mats (Pascal et al., 2017) are still exposed to higher concentrations of Hg in the substrate. Even if the dominant form of Hg in organisms is likely inorganic Hg, methylation of Hg could occur in their gut too (Rudd et al., 1980; Li et al., 2018), which could induce a greater biomagnification potential in Bouillante, since MeHg can enter in the cytoplasm and is thus more easily transferred through the food chain (Baeyens et al., 2003). However, it has been shown recently that demethylation can also occur in fish guts (Li et al., 2018), and further studies should therefore focus more closely on Hg speciation in Bouillante’s food webs.

**Figure 4.**
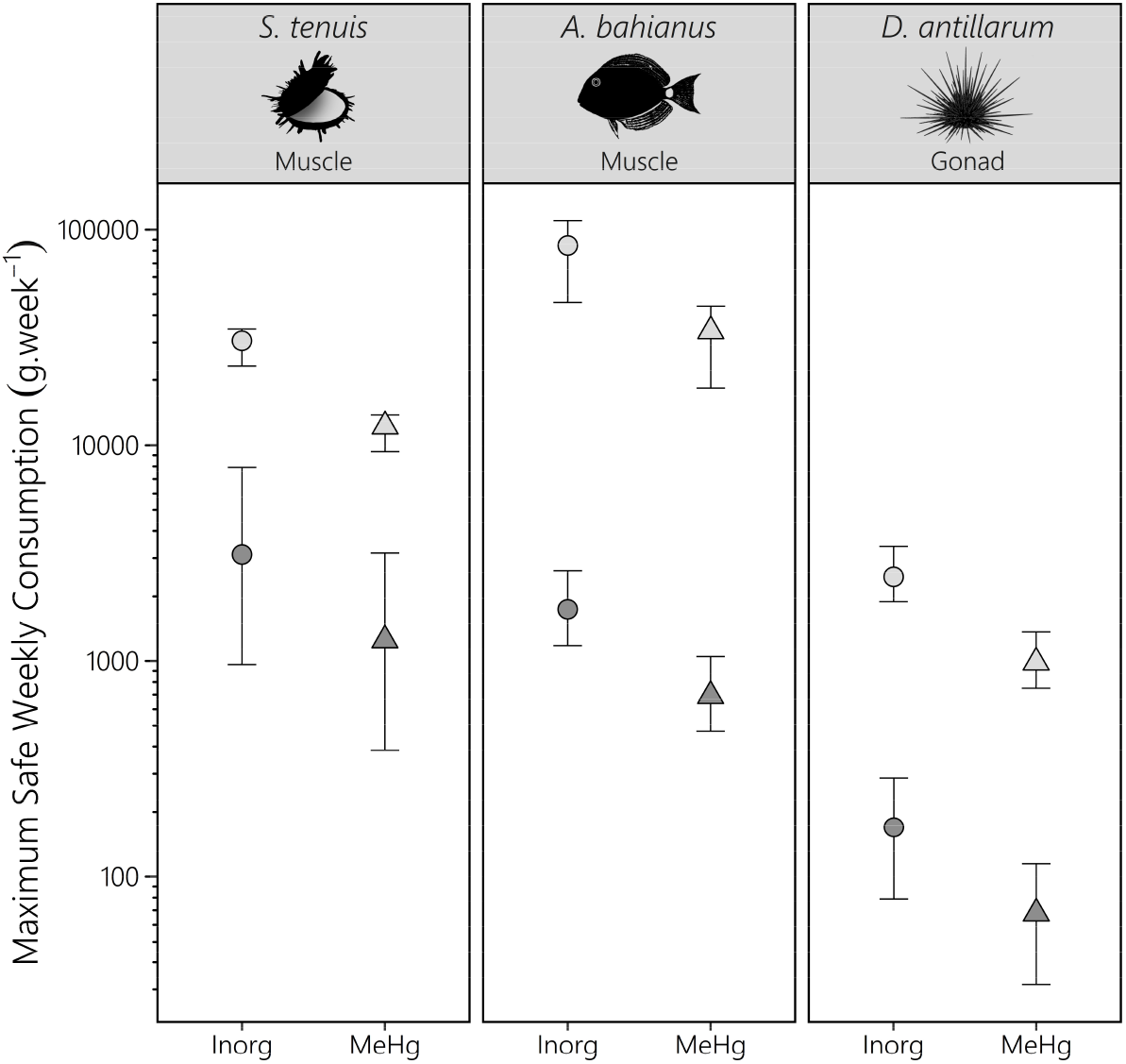
Maximum Safe Weekly Consumption (MSWC) quantities according to the FAO guidelines for the three species potentially eaten by humans. The different colours represent the different sites (light grey = Control Site, dark grey = Bouillante) while the different shapes represent the different hypotheses concerning the type of mercury bioavailable in the Bouillante Bay (dot = inorganic Hg; triangle = organic Hg). The vertical bars represent the values obtained after calculation of the MSCW on the limits of the 95% bootstrapped confidence interval of the Hg concentration (iterations = 1,000,000).

**Figure 5.**
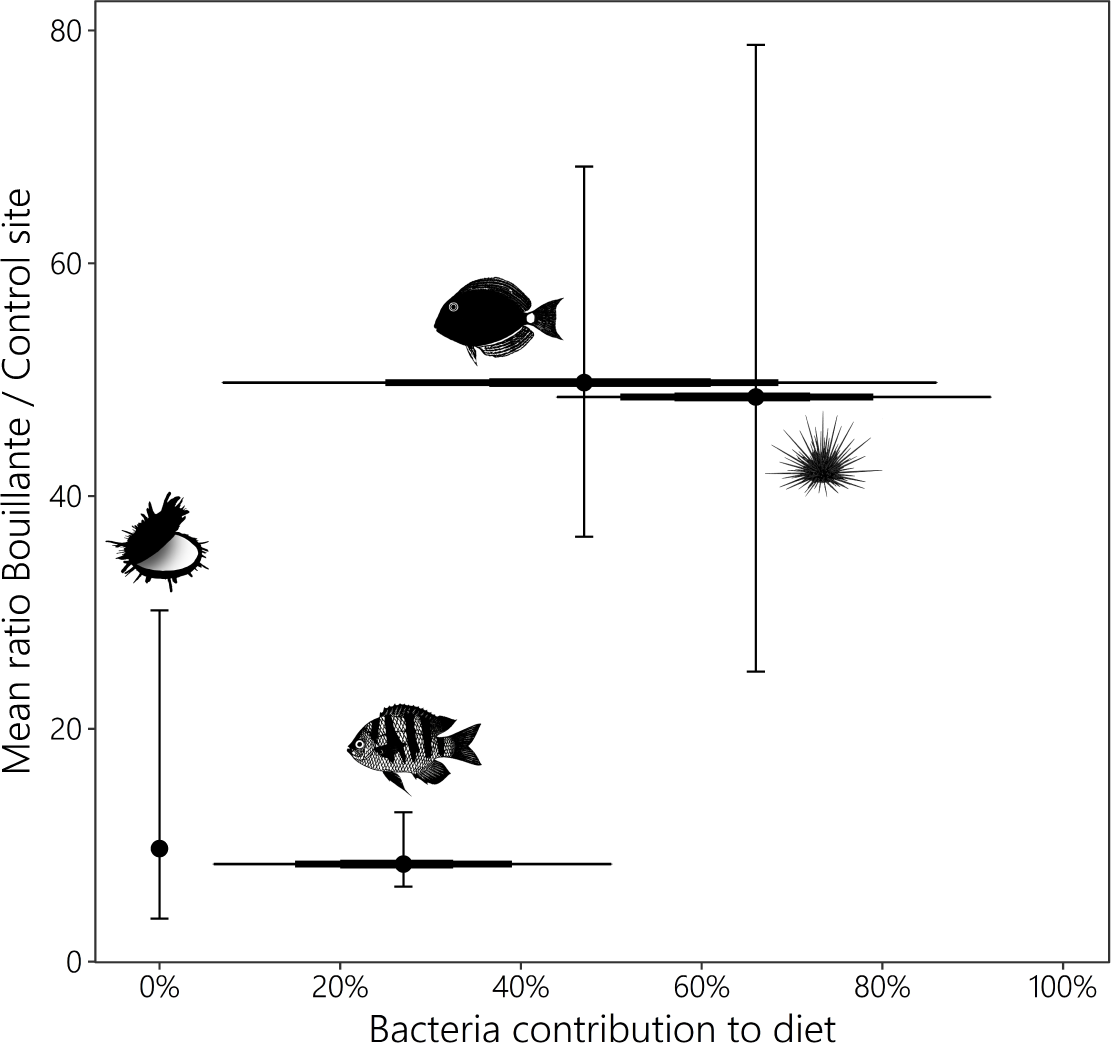
Mean ratio between the Hg concentration in muscles collected in Bouillante and the Control Site compared with the bacteria contribution to diet found for other individuals of the same species by Pascal et al. (2017) in the Bouillante Bay. The three different thickness represent respectively the 25%, 75% and 95% credibility intervals while the vertical bars represent the 95% confidence interval around the mean ratio of Hg concentration (iterations = 1,000,000).

The main issue however probably concerns the health of Bouillante marine organisms themselves, since some forms of inorganic Hg, such as HgCl_2_, are even more nephrotoxic than MeHg (Zalups, 2000). For example, it was shown that only 4 days after the injection of 0.6 μg.g^-1^ of HgCl_2_, a concentration 88 (*A. saxatilis*) and 128 times (*A. bahianus*) lower than in the liver of Bouillante fishes, severe disorganisation and degradation were already observed in *Gymnotus carapo* liver tissues (Vergilio et al., 2012). More generally, seeing concentrations in livers, fishes of Bouillante probably also contains very high levels of Hg in their kidneys, gills and gut, which would occasion breathing problems, endocrine disruptions, starvation, acceleration of metabolism, and severe stress (Zulkipli et al., 2021). Moreover, our results reveal that sea urchins gonads contain abnormally high levels of mercury in Bouillante but a 20-30mn exposure to a 2.7-3.3 μg.L^-1^ of HgCl_2_ has been shown to completely inhibit sperm fertilization capacities in another sea urchin species (Eissa, 2007). Globally, the strong affinity of Hg with thiol (–SH) groups found in many enzymes disrupts all kinds of biological processes and has no known biological function in any living being (Liu et al., 2012). Therefore, short-term advantages (e.g. abundant food supply, silica from geothermal fluid favouring sponge growth) described in Pascal et al. (2017), which results in a higher abundance of some opportunistic species near the discharge channel, might be counterbalanced by long-term harmful effects on their health. This result contradicts all previous impact assessments on the Bouillante geothermal plant, which classified the environmental effect of the release of discharge water into the sea as limited (PARETO – IMPACTMER, 2009) or negligible (ADEME – BRGM, 2004; ANTEA, 2005).

## Conclusion

By comparing Hg concentrations in animals and sediment samples from the Bouillante Bay to a Control Site and prospection studies preceding the construction of the geothermal power plant, it appears that the discharge of hydrothermal waters into the sea can be clearly linked with the significant Hg contamination we detected. Two pathways of Hg integration into Bouillante’s food webs through the low trophic levels animals monitored in this study were emphasised. First, greater concentrations in filter-feeders not consuming bacterial mats (shown through isotopic analysis; Pascal et al., 2017) and liver/muscle ratios in fishes indicate that discharge waters released at high rates by the power plant probably enhance the exposure of marine organisms to dissolved inorganic Hg (i.e. diffusive pathway). Second, opportunistic animals taking advantage as a new food source of the sulphur-bacterial mats (Pascal et al., 2017) growing in the channel thanks to the abundant sulphur supply accumulate elevated and potentially harmful levels of Hg (i.e. dietary pathway). Indeed, bacterial mats efficiently entrap Hg in their EPS as a detoxification mechanism, and thus accumulate very high levels of Hg. Generally, we demonstrate for the first time the role of sulphur-oxidising bacteria in the transfer of inorganic Hg, which enlightens our understanding of vent fields as natural sources of Hg.

## Acknowledgements

This work was supported by the project ECONAT “Axe 1 - Ressources Marines et Littorales : Qualité et Eco-valorisation” (QUALIDRIS), funded by the Contrat de Plan Etat-Région (CPER), the CNRS and the European Regional Development Fund (FEDER). Authors warmly thank Christine Dupuy for her key role in the coordination of these projects. We also thank Sebastien Cordonnier for his assistance in the field. The authors declare that they have no known competing financial interests or personal relationships that could have appeared to influence the work reported in this paper.

## References

1. ADEME-BRGM, 2004. La géothermie. Collection « Les enjeux des géosciences », BRGM.

2. Adeyanju, E., & Okeke, C. A. (2019). Exposure effect to cement dust pollution: a mini review. SN Applied Sciences, 1(12), 1–17.

3. Amos, H. M., Jacob, D. J., Streets, D. G., & Sunderland, E. M. (2013). Legacy impacts of all-time anthropogenic emissions on the global mercury cycle. Global Biogeochemical Cycles, 27(2), 410–421.

4. ANTEA (2005). Etude d’impact de l’unité Bouillante 2. Rapport A31753/C, CFG Services, Orléans

5. Baeyens, W., Leermakers, M., Papina, T., Saprykin, A., Brion, N., Noyen, J., De Gieter, M., Elskens, M., & Goeyens, L. (2003). Bioconcentration and biomagnification of mercury and methylmercury in North Sea and Scheldt Estuary fish. Archives of Environmental Contamination and Toxicology, 45(4), 498–508.

6. Bagnato, E., Aiuppa, A., Parello, F., D’alessandro, W., Allard, P., & Calabrese, S. (2009). Mercury concentration, speciation and budget in volcanic aquifers: Italy and Guadeloupe (Lesser Antilles). Journal of Volcanology and Geothermal Research, 179(1-2), 96–106.

7. Balogh, S. J., Tsui, M. T. K., Blum, J. D., Matsuyama, A., Woerndle, G. E., Yano, S., & Tada, (2015). Tracking the fate of mercury in the fish and bottom sediments of Minamata Bay, Japan, using stable mercury isotopes. Environmental science & technology, 49(9), 5399–5406.

8. Bargagli, R., Monaci, F., Sanchez-Hernandez, J. C., & Cateni, D. (1998). Biomagnification of mercury in an Antarctic marine coastal food web. Marine Ecology Progress Series, 169, 65–76.

9. Barlow, S., Bolger, M., DiNovi, M., Renwick, A., Street, D., & Schlatter J. (2007). Safety evaluation of certain food additives and contaminants – Methylmercury. World Health Organization (FAO JECFA), Geneva, 58, 269–315.

10. Batista, D., Muricy, G., Rocha, R. C., & Miekeley, N. F. (2014). Marine sponges with contrasting life histories can be complementary biomonitors of heavy metal pollution in coastal ecosystems. Environmental Science and Pollution Research, 21(9), 5785–5794.

11. Beaulieu, S.E., & Szafranski, K. (2020). InterRidge Global Database of Active Submarine Hydrothermal Vent Fields, Version 3.4. World Wide Web electronic publication available from: http://vents-data.interridge.org. Accessed 2022-01-22.

12. Benjamini, Y., & Hochberg, Y. (1995). Controlling the false discovery rate: a practical and powerful approach to multiple testing. Journal of the Royal statistical society: series B (Methodological), 57(1), 289–300.

13. Bjerregaard, P., & Møller, L. M. (2021). Exposure to methylmercury and inorganic mercury in the food does not lead to trophic magnification in the sea star Asterias rubens. Environmental Pollution, 285, 117401.

14. Bosch, A. C., O’Neill, B., Sigge, G. O., Kerwath, S. E., & Hoffman, L. C. (2015). Heavy metals in marine fish meat and consumer health: a review. Journal of the Science of Food and Agriculture, 96(1), 32–48.

15. Caraïbes Environnement (2011). Etude d’impact sur l’environnement.

16. Cardigos, F., Colaço, A., Dando, P. R., Ávila, S. P., Sarradin, P. M., Tempera, F., … & Santos, R. S. (2005). Shallow water hydrothermal vent field fluids and communities of the D. João de Castro Seamount (Azores). Chemical Geology, 224(1-3), 153–168.

17. Chapman, M. R., & Kramer, D. L. (2000). Environmental Biology of Fishes, 57(1), 11–24.

18. Chou, C. Y., Lin, Y. C., Chang, C. L., & Chen, C. H. (2006). On the bootstrap confidence intervals of the process incapability index Cpp. Reliability Engineering & System Safety, 91(4), 452–459.

19. Chouvelon, T., Warnau, M., Churlaud, C., & Bustamante, P. (2009). Hg concentrations and related risk assessment in coral reef crustaceans, molluscs and fish from New Caledonia. Environmental Pollution, 157(1), 331–340.

20. Cizdziel, J. V., Hinners, T. A., Pollard, J. E., Heithmar, E. M., & Cross, C. L. (2002). Mercury concentrations in fish from Lake Mead, USA, related to fish size, condition, trophic level, location, and consumption risk. Archives of Environmental Contamination and Toxicology, 43(3), 0309–0317.

21. Colombet, L. N. (2021). Inventaire National du Patrimoine Naturel. Observations naturalistes des Amis de BioObs. Version 1.1. UMS PatriNat (OFB-CNRS-MNHN), Paris. Occurrence dataset https://doi.org/10.15468/ldch7a accessed via GBIF.org on 2022-01-07. https://www.gbif.org/occurrence/3019231290

22. Coppard, S. E., & Campbell, A. C. (2004). Taxonomic significance of spine morphology in the echinoid genera Diadema and Echinothrix. Invertebrate Biology, 123(4), 357–371.

23. Crocker, L. A., & Reiswig, H. M. (1981). Host specificity in sponge-encrusting zoanthidea (Anthozoa: Zoantharia) of Barbados, West Indies. Marine Biology, 65(3), 231–236.

24. Cruz, K. A. (2014). Extracellular polysaccharides production by bacteria as a mechanism of mercury tolerance (Doctoral dissertation, Rutgers University-Graduate School-New Brunswick).

25. Cudney-Bueno, R., & Rowell, K. (2008). Establishing a baseline for management of the rock scallop, Spondylus calcifer (Carpenter 1857): Growth and reproduction in the upper Gulf of California, Mexico. Journal of Shellfish Research, 27(4), 625–632.

26. De Mestre, C., Maher, W., Roberts, D., Broad, A., Krikowa, F., & Davis, A. R. (2012). Sponges as sentinels: patterns of spatial and intra-individual variation in trace metal concentration. Marine pollution bulletin, 64(1), 80–89.

27. De Zoysa, H. K. S., Jinadasa, B. K. K. K., Edirisinghe, E. M. R. K. B., & Jayasinghe, G. D. T. M. (2018). The association of test diameter and gonad weight with some toxic trace metals level in black sea urchin (Stomopneustes variolaris). Agriculture & Food Security, 7(1), 1–12.

28. Demarcq, F., Vernier, R., & Sanjuan, B. (2014, April). Situation and perspectives of the Bouillante geothermal power plant in Guadeloupe, French West Indies. In Deep Geothermal Days.

29. Denton, G. R. W., & Burdon-Jones, C. (1986). Trace metals in fish from the Great Barrier Reef. Marine Pollution Bulletin, 17(5), 201–209.

30. Denton, G. R. W., Concepcion, L. P., Wood, H. R., & Morrison, R. J. (2006). Trace metals in marine organisms from four harbours in Guam. Marine pollution bulletin, 52(12), 1784–1804.

31. Denton, G. R., Wood, H. R., Concepcion, L. P., Siegrist, H. G., Eflin, V. S., Narcis, D. K., & Pangelinan, G. T. (1997). Analysis of in-place contaminants in marine sediments from four harbor locations on Guam. Water and Environmental Research Institute of the Western Pacific, University of Guam.

32. Dixit, C. (2014). Etude physico-chimique des fluides produits par la centrale géothermique de Bouillante (Guadeloupe) et des dépôts susceptibles de se former au cours de leur refroidissement (Doctoral dissertation, Antilles-Guyane).

33. Ebert, T. A., Russell, M. P., Gamba, G., & Bodnar, A. (2008). Growth, survival, and longevity estimates for the rock-boring sea urchin Echinometra lucunter lucunter (Echinodermata, Echinoidea) in Bermuda. Bulletin of Marine Science, 82(3), 381–403.

34. Eissa, S. H. H. (2007). The toxicity of mercury on the sea urchin gametes and embryos. Egyptian Journal of Experimental Biology, 3, 37–42.

35. Elliott, J. E., Kirk, D. A., Elliott, K. H., Dorzinsky, J., Lee, S., Inzunza, E. R., … & Shaw, P. (2015). Mercury in forage fish from Mexico and Central America: implications for fish-eating birds. Archives of environmental contamination and toxicology, 69(4), 375–389.

36. Erwin, P. M., & Thacker, R. W. (2007). Incidence and identity of photosynthetic symbionts in Caribbean coral reef sponge assemblages. Journal of the Marine Biological Association of the United Kingdom, 87(6), 1683–1692.

37. Fabriol R., Ouzounian G. (1985) - Prospection géothermique des zones de Bouillante et de la Soufrière (Guadeloupe), Modèle hydrogéochimique. Rapport BRGM 85 SGN 433 GTH, 29 p.

38. Fabriol, R., & Hazan, M. (1984). Prospection des anomalies de mercure dans les sols. Prospection géothemique de la région de Bouillante – Vieux-habitants Guadeloupe. Bureau de recherche géologique et Minières, Service géologique et national.

39. Feeley, M., Barraj, L., Bellinger, D.C., Bronson, R., Guérin, T., Larsen, J.C, Lo, M.-T. & Slob, W. (2011). Safety evaluation of certain food additives and contaminants – Mercury (addendum). World Health Organization (FAO JECFA), Geneva, 63(8), 605–684.

40. Frederick, P., Axelrad, D., Atkson, T., & Pollman, C. (2005). Contaminants research and policy: the Everglades mercury story. National Wetlands Newsletter, 27(1), 3–6.

41. Fujiki, M., & Tajima, S. (1992). The pollution of Minamata Bay by mercury. Water Science and Technology, 25(11), 133–140.

42. Gadalia, A., Bouchot, V., Calcagno, P., Caritg, S., Courrioux, G., Darnet, M., Jacob, T., Labeau, Y., Taïlamé, A. L., Terrier, M., Thinon, I., & Vittecoq, B. (2019, March). Multimodal geothermal exploration in the Lesser Antilles Arc at the Lamentin lowland (Martinique). In IOP Conference Series: Earth and Environmental Science (Vol. 249, No. 1, p. 012001). IOP Publishing.

43. Géothermie Bouillante (2018). Demande d’Autorisation d’Ouverture de Travaux miniers pour la réalisation de nouveaux forages – Etude d’Impact. Concession géothermique de Bouillante.

44. Gionfriddo, C. M., Tate, M. T., Wick, R. R., Schultz, M. B., Zemla, A., Thelen, M. P., … & Moreau, J. W. (2016). Microbial mercury methylation in Antarctic sea ice. Nature microbiology, 1(10), 16127.

45. Gobert, B., & Reynal, L. (2002). Les ressources démersales des Antilles et leur exploitation. La pêche aux Antilles: Martinique et Guadeloupe, IRD Editions, pp. 49b65.

46. Gremyachikh, V. A., Lozhkina, R. A., & Komov, V. T. (2019). Spatial-temporal variability of mercury content in the river perch Perca fluviatilis Linnaeus, 1758 (Perciformes: Percidae) of the Rybinsk Reservoir at the turn of the XX–XXI centuries. Ecosystem Transformation, (2).

47. Grisolía, J. M., López, F., & de Dios Ortuzar, J. (2012). Sea urchin: From plague to market opportunity. Food quality and preference, 25(1), 46–56.

48. Guézennec, J., Moppert, X., Raguénès, G., Richert, L., Costa, B., & Simon-Colin, C. (2011). Microbial mats in French Polynesia and their biotechnological applications. Process Biochemistry, 46(1), 16–22.

49. Han, R., Zhou, B., Huang, Y., Lu, X., Li, S., & Li, N. (2020). Bibliometric overview of research trends on heavy metal health risks and impacts in 1989–2018. Journal of Cleaner Production, 276, 123249.

50. Hansen, M., & Perner, M. (2016). Hydrogenase gene distribution and H2 consumption ability within the Thiomicrospira lineage. Frontiers in microbiology, 7, 99.

51. Hartwell, S. I., Apeti, D. A., Pait, A. S., Mason, A. L., & Robinson, C. M. (2017). An analysis of chemical contaminants in sediments and fish from Cocos Lagoon, Guam.

52. Hawkins, J. P., Roberts, C. M., Gell, F. R., & Dytham, C. (2007). Effects of trap fishing on reef fish communities. Aquatic Conservation: Marine and Freshwater Ecosystems, 17(2), 111–132.

53. Hsu-Kim, H., Kucharzyk, K. H., Zhang, T., & Deshusses, M. A. (2013). Mechanisms regulating mercury bioavailability for methylating microorganisms in the aquatic environment: a critical review. Environmental science & technology, 47(6), 2441–2456.

54. Jiskra, M., Heimbürger-Boavida, L. E., Desgranges, M. M., Petrova, M. V., Dufour, A., Ferreira-Araujo, B., … & Sonke, J. E. (2021). Mercury stable isotopes constrain atmospheric sources to the ocean. Nature, 597(7878), 678–682.

55. Keshavarz, M., & Jahromi, M. S. (2017). Effects of primary sex ratio on operational sex ratio in sea urchin, Echinometra mathaei. Pakistan Journal of Zoology, 49(4), 1373–1381.

56. Kleint, C., Pichler, T., & Koschinsky, A. (2017). Geochemical characteristics, speciation and size-fractionation of iron (Fe) in two marine shallow-water hydrothermal systems, Dominica, Lesser Antilles. Chemical Geology, 454, 44–53.

57. Lachassagne, P., Maréchal, J. C., & Sanjuan, B. (2009). Hydrogeological model of a high-energy geothermal field (Bouillante area, Guadeloupe, French West Indies). Hydrogeology Journal, 17(7), 1589–1606.

58. Lacoue-Labarthe, T., Warnau, M., Beaugeard, L., & Pascal, P. Y. (2016). Trophic transfer of radioisotopes in Mediterranean sponges through bacteria consumption. Chemosphere, 144, 1885–1892.

59. Lefebvre, D. D., Kelly, D., & Budd, K. (2007). Biotransformation of Hg (II) by cyanobacteria. Applied and Environmental Microbiology, 73(1), 243–249.

60. Lewis, S. M. (1982). Sponge-zoanthid associations: Functional interactions. Smithsonian Contributions to the Marine Sciences, 12, 465–474.

61. Li, H., Lin, X., Zhao, J., Cui, L., Wang, L., Gao, Y., … & Li, Y. F. (2018). Intestinal methylation and demethylation of mercury. Bulletin of environmental contamination and toxicology, 102(5), 597–604.

62. Liu, G., Cai, Y., O’Driscoll, N., Feng, X., & Jiang, G. (2012). Overview of mercury in the environment. Environmental chemistry and toxicology of mercury, 1–12.

63. Macieira, R. M., & Joyeux, J. C. (2008). Short communication Length–weight relationships for rockpool fishes in Brazil. J. Appl. Ichthyol, 1, 2.

64. Manullang, C. Y., Cordova, M. R., Purbonegoro, T., Soamole, A., & Rehalat, I. (2020, December). Mercury concentrations in Kayeli Bay, Buru Island of Indonesia: The update of possible effect of land-based gold mining. In IOP Conference Series: Earth and Environmental Science (Vol. 618, No. 1, p. 012023). IOP Publishing.

65. Martins, I., Costa, V., Porteiro, F. M., Colaço, A., & Santos, R. S. (2006). Mercury concentrations in fish species caught at Mid-Atlantic Ridge hydrothermal vent fields. Marine Ecology Progress Series, 320, 253–258.

66. Martins, I., Costa, V., Porteiro, F., Cravo, A., & Santos, R. S. (2001). Mercury concentrations in invertebrates from Mid-Atlantic Ridge hydrothermal vent fields. Journal of the Marine Biological Association of the United Kingdom, 81(6), 913–915.

67. Mas, A., Guisseau, D., Mas, P. P., Beaufort, D., Genter, A., Sanjuan, B., & Girard, J. P. (2006). Clay minerals related to the hydrothermal activity of the Bouillante geothermal field (Guadeloupe). Journal of Volcanology and Geothermal research, 158(3-4), 380–400.

68. Maury-Brachet R., Durrieu, G., Dominique, Y., & Boudou, A. (2006). Mercury distribution in fish organs and food regimes: Significant relationships from twelve species collected in French Guiana (Amazonian basin). Science of the Total Environment, 368(1), 262–270.

69. Medeiros, P. R., Grempel, R. G., Souza, A. T., Ilarri, M. I., & Sampaio, C. L. S. (2007). Effects of recreational activities on the fish assemblage structure in a northeastern Brazilian reef. Pan-American Journal of Aquatic Sciences, 2(3), 288–300.

70. Micheline, G., Rachida, C., Céline, M., Gaby, K., Rachid, A., & Petru, J. (2019). Levels of Pb, Cd, Hg and As in fishery products from the Eastern Mediterranean and human health risk assessment due to their consumption. International Journal of Environmental Research, 13(3), 443–455.

71. Miller, J. W., VanDerwalker, J. G., & Waller, R. A. (1971). Scientists-in-the-sea. US Department of the Interior. Washington DC.

72. Mir, J., Martínez-Alonso, M., Esteve, I., & Guerrero, R. (1991). Vertical stratification and microbial assemblage of a microbial mat in the Ebro Delta (Spain). FEMS Microbiology Letters, 86(1), 59–68.

73. Morrison, R. J., Peshut, P. J., West, R. J., & Lasorsa, B. K. (2015). Mercury (Hg) speciation in coral reef systems of remote Oceania: Implications for the artisanal fisheries of Tutuila, Samoa Islands. Marine pollution bulletin, 96(1-2), 41–56.

74. National Center for Biotechnology Information (NCBI)[Internet]. Bethesda (MD): National Library of Medicine (US), National Center for Biotechnology Information; [1988] – [cited 2022 Apr 21]. Available from: https://www.ncbi.nlm.nih.gov/

75. Pan, K., & Wang, W. X. (2011). Mercury accumulation in marine bivalves: influences of biodynamics and feeding niche. Environmental pollution, 159(10), 2500–2506.

76. Pan, K., Lee, O. O., Qian, P. Y., & Wang, W. X. (2011). Sponges and sediments as monitoring tools of metal contamination in the eastern coast of the Red Sea, Saudi Arabia. Marine pollution bulletin, 62(5), 1140–1146.

77. PARETO-IMPACTMER (2009) Rejets en mer de la centrale géothermique de Bouillante (Unité 1 et 2): compléments à l’étude d’impact de 2005, étude des biocénoses marines, CFG Service / Géothermie de Bouillante, Orléans.

78. Parks, J. M., Johs, A., Podar, M., Bridou, R., Hurt, R. A., Smith, S. D., Tomaniceck, S. J., Quian, Y., Brown, S. D., Brandt, C. C., Palumbo, A. V., Smith, J. C., Wall, J. D., Elias, D. A., & Liang, L. (2013). The genetic basis for bacterial mercury methylation. Science, 339(6125), 1332–1335.

79. Pascal, P. Y., Dubois, S. F., Goffette, A., & Lepoint, G. (2017). Influences of geothermal sulfur bacteria on a tropical coastal food web. Marine Ecology Progress Series, 578, 73–85.

80. Pawlik, J. R., McMurray, S. E., Erwin, P., & Zea, S. (2015). No evidence for food limitation of Caribbean reef sponges: reply to Slattery & Lesser (2015). Marine Ecology Progress Series, 527, 281–284.

81. Perez, T. (2004). In situ comparative study of several Mediterranean sponges as potential biomonitors of heavy metals. BMIB-Bollettino dei Musei e degli Istituti Biologici, 68.

82. Pickhardt, P. C., Stepanova, M., & Fisher, N. S. (2006). Contrasting uptake routes and tissue distributions of inorganic and methylmercury in mosquitofish (Gambusia affinis) and redear sunfish (Lepomis microlophus). Environmental Toxicology and Chemistry: An International Journal, 25(8), 2132–2142.

83. Podar, M., Gilmour, C. C., Brandt, C. C., Soren, A., Brown, S. D., Crable, B. R., … & Elias, D. (2015). Global prevalence and distribution of genes and microorganisms involved in mercury methylation. Science advances, 1(9), e1500675.

84. Robertson, D., Ackerman, J., Choat, J., Posada, J., & Pitt, J. (2005). Ocean surgeonfish Acanthurus Bahianus. i. the geography of Demography. Marine Ecology Progress Series, 295, 229–244. doi:10.3354/meps295229

85. Rocha, V. P., Silveira, I. D. O., & Matthews-Cascon, H. (2015). Brazilian Spondylidae: a brief discussion about variation of shell ornamentation in the Northeastern species.

86. Roos-Barraclough, F., Givelet, N., Martinez-Cortizas, A., Goodsite, M. E., Biester, H., & Shotyk, W. (2002). An analytical protocol for the determination of total mercury concentrations in solid peat samples. Science of the total environment, 292(1-2), 129–139.

87. Rudd, J. W., Furutani, A. K. I. R. A., & Turner, M. A. (1980). Mercury methylation by fish intestinal contents. Applied and Environmental Microbiology, 40(4), 777–782.

88. Rumbold, D. G., Evans, D. W., Niemczyk, S., Fink, L. E., Laine, K. A., Howard, N., … & Zucker, M. (2011). Source identification of Florida Bay’s methylmercury problem: mainland runoff versus atmospheric deposition and in situ production. Estuaries and coasts, 34(3), 494–513.

89. Rumbold, D. G., Lienhardt, C. T., & Parsons, M. L. (2018). Mercury biomagnification through a coral reef ecosystem. Archives of environmental contamination and toxicology, 75(1), 121–133.

90. Sanjuan, B., & Brach, M. (1997). Etude hydrogéochimique du champ géothermique de Bouillante (Guadeloupe). Rapport BRGM/R-39880-FR.

91. Sanjuan, B., Lasne, E., & Brach, M. (2000). Bouillante geothermal field (Guadeloupe, West Indies): Geochemical monitoring during a thermal stimulation operation. Twenty-Fifth Workshop on Geothermal Reservoir Engineering Stanford University, Stanford, California, United States. pp.215–222.

92. Sanjuan, B., Le Nindre, Y. M., Menjoz, A., Sbai, A., Brach, M., & Lasne, E. (2004). Travaux de recherche liés au développement du champ géothermique de Bouillante (Guadeloupe). Rapport BRGM/RP-53136-FR.

93. Schaefer, J. K., & Morel, F. M. (2009). High methylation rates of mercury bound to cysteine by Geobacter sulfurreducens. Nature geoscience, 2(2), 123–126.

94. Scheuhammer, A., Braune, B., Chan, H. M., Frouin, H., Krey, A., Letcher, R., … & Wayland, M. (2015). Recent progress on our understanding of the biological effects of mercury in fish and wildlife in the Canadian Arctic. Science of the Total Environment, 509, 91–103.

95. Schlieman, C. (2020). Regional differences driving organic matter and trace metal signatures reflected in temperate reef bivalve communities on the South Island, New Zealand (Doctoral dissertation, University of Otago).

96. Silvano, R. A., & Begossi, A. (2012). Fishermen’s local ecological knowledge on Southeastern Brazilian coastal fishes: contributions to research, conservation, and management. Neotropical Ichthyology, 10(1), 133–147.

97. Stolz, J. F. (1985). The microbial community at Laguna Figueroa, Baja California Mexico: from miles to microns. Origins of life and evolution of the biosphere, 15(4), 347–352.

98. Szkoda, J., Durkalec, M., Nawrocka, A., & Michalski, M. (2015). Mercury concentration in bivalve molluscs. Bull Vet Inst Pulawy, 59, 357–360.

99. Takai, K., Nealson, K. H., & Horikoshi, K. (2004). Hydrogenimonas thermophila gen. nov., sp. nov., a novel thermophilic, hydrogen-oxidizing chemolithoautotroph within the ε-Proteobacteria, isolated from a black smoker in a Central Indian Ridge hydrothermal field. International Journal of Systematic and Evolutionary Microbiology, 54(1), 25–32.

100. Tan, G., Sun, W., Xu, Y., Wang, H., & Xu, N. (2016). Sorption of mercury (II) and atrazine by biochar, modified biochars and biochar based activated carbon in aqueous solution. Bioresource technology, 211, 727–735.

101. Trager, G. C., & DeNiro, M. J. (1990). Chemoautotrophic sulfur bacteria as a food source for mollusks at intertidal hydrothermal vents: evidence from stable isotopes. The Veliger, 33(4), 359–362.

102. Uihlein A., (2018). JRC Geothermal Power Plant Dataset – Documentation EUR 29446 EN. Publications Office of the European Union, Luxembourg.

103. Vasconcelos, P., Carvalho, A. N., Moura, P., Ramos, J., & Gaspar, M. B. (2018). First record of Acanthurus monroviae (Osteichthyes: Perciformes: Acanthuridae) in southern Portugal, with notes on its recent distributional spread in the northeastern Atlantic and Mediterranean. Marine Biodiversity, 48(3), 1673–1681.

104. Vergilio, C. S., Carvalho, C. E. V., & DE MELO, E. J. T. (2012). Accumulation and histopathological effects of mercury chloride after acute exposure in tropical fish Gymnotus carapo.

105. Vidal, D. E., & Horne, A. J. (2003). Inheritance of mercury tolerance in the aquatic oligochaete Tubifex tubifex. Environmental Toxicology and Chemistry: An International Journal, 22(9), 2130–2135.

106. Vieira, J. C. S., Braga, C. P., de Oliveira, G., de Lima Leite, A., de Queiroz, J. V., Cavecci, B., … & de Magalhães Padilha, P. (2017). Identification of protein biomarkers of mercury toxicity in fish. Environmental Chemistry Letters, 15(4), 717–724.

107. Volesky, B. (1990). Removal and recovery of heavy metals by biosorption. Biosorption of heavy metals, 7–43.

108. Walpole, S. C., Prieto-Merino, D., Edwards, P., Cleland, J., Stevens, G., & Roberts, I. (2012). The weight of nations: an estimation of adult human biomass. BMC public health, 12(1), 1–6.

109. Wang, W. X., & Wong, R. S. (2003). Bioaccumulation kinetics and exposure pathways of inorganic mercury and methylmercury in a marine fish, the sweetlips Plectorhinchus gibbosus. Marine Ecology Progress Series, 261, 257–268.

110. Warnau, M., Ledent, G., Temara, A., Bouquegneau, J. M., Jangoux, M., & Dubois, P. (1995). Heavy metals in Posidonia oceanica and Paracentrotus lividus from seagrass beds of the north-western Mediterranean. Science of the total environment, 171(1-3), 95–99.

111. Weis, J. S. (2002). Tolerance to environmental contaminants in the mummichog, Fundulus heteroclitus. Human and Ecological Risk Assessment, 8(5), 933–953.

112. Weisz, J. B., Lindquist, N., & Martens, C. S. (2008). Do associated microbial abundances impact marine demosponge pumping rates and tissue densities?. Oecologia, 155(2), 367–376.

113. Zalups, R. K. (2000). Molecular interactions with mercury in the kidney. Pharmacological reviews, 52(1), 113–144.

114. Zulkipli, S. Z., Liew, H. J., Ando, M., Lim, L. S., Wang, M., Sung, Y. Y., & Mok, W. J. (2021). A review of mercury pathological effects on organs specific of fishes. Environmental Pollutants and Bioavailability, 33(1), 76–87.

