## Supplementary Tables for "Hydrothermal sulphur bacteria enhance mercury availability for coastal marine organisms"

| Species /  Type | Tissue | Control (mean) | Bouillante (mean) | Mean ratio B/C |
| --- | --- | --- | --- | --- |
| Surface sediment | Ø | 0.024 μg.g^-1^  95% CI [0.023, 0.025] | 0.326 μg.g^-1^ | × 13.63 |
| Bacterial mat | Ø | Ø | 13.01 μg.g^-1^ | Ø |
| *Iotrochota birotulata* | Ø | 0.066 μg.g^-1^  95% CI [0.060, 0.072] | 0.176 μg.g^-1^  95% CI [0.163, 0.188] | × 2.64  95% CI [2.37, 2.96] |
| *Aplysina fistularis* | Ø | 0.477 μg.g^-1^  95% CI [0.416, 0.519] | 1.164 μg.g^-1^  95% CI [1.094, 1.220] | × 2.44  95% CI [2.20, 2.81] |
| *Spondylus tenuis* | Muscle | 0.036 μg.g^-1^  95% CI [0.032, 0.047] | 0.353 μg.g^-1^  95% CI [0.139, 1.143] | × 9.71  95% CI [3.69, 30.02] |
| *Abudefduf saxatilis* | Muscle | 0.043 μg.g^-1^  95% CI [0.034, 0.049] | 0.361 μg.g^-1^  95% CI [0.293, 0.528] | × 8.38  95% CI [6.43, 12.78] |
| *Acanthurus bahianus* | Muscle | 0.013 μg.g^-1^  95% CI [0.010, 0.024] | 0.637 μg.g^-1^  95% CI [0.420, 0.935] | × 48.51  95% CI [24.96, 78.77] |
| *Diadema antillarum* | Muscle | 0.032 μg.g^-1^  95% CI [0.028, 0.039] | 1.601 μg.g^-1^  95% CI [1.253, 2.108] | × 49.75  95% CI [36.50, 68.34] |
| *Diadema antillarum* | Gonad | 0.447 μg.g^-1^  95% CI [0.324, 0.587] | 6.498 μg.g^-1^  95% CI [3.84, 13.93] | × 14.52  95% CI [8.01, 34.80] |
| *Abudefduf saxatilis* | Liver | 0.084 μg.g^-1^  95% CI [0.038, 0.221] | 52.88 μg.g^-1^  95% CI [30.60, 79.89] | × 626.82  95% CI [191.24, 1589.50] |
| *Acanthurus bahianus* | Liver | 1.879 μg.g^-1^  95% CI [0.866, 3.885] | 77.22 μg.g^-1^  95% CI [45.70, 109.9] | × 41.09  95% CI [16.22, 99.77] |

**Table S1** – Concentrations of mercury in Bouillante and the Control Site, as well as mean ratios Bouillante / Control, with their robust 95% CI (calculated after 1,000,000 bootstrap iterations). Concentrations were calculated based on dry weight.

| Species | Tissue | Test | X | df | *p* | N | Predicted difference (B – C) | Effect size |
| --- | --- | --- | --- | --- | --- | --- | --- | --- |
| *Iotrochota birotulata* | Ø | Student (Welch correction) | t = 14.5 | 12.2 | < .001 | 20 | 0.11 μg.g^-1^  95% CI [0.09, 0.13] | d = 6.49 (large)  95% CI [4.14, 8.84] |
| *Aplysina fistularis* | Ø | Student | t = 15.8 | 18 | < .001 | 20 | 0.69 μg.g^-1^  95% CI [0.60, 0.78] | d = 7.08 (large)  95% CI [4.55, 9.61] |
| *Spondylus tenuis* | Muscle | Wilcoxon | W = 99 | Ø | < .001 | 20 | 0.12 μg.g^-1^  95% CI [0.08, 0.20] | r = 0.83 (large)  95% CI [0.71, 0.85] |
| *Abudefduf saxatilis* | Muscle | Wilcoxon | W = 100 | Ø | < .001 | 20 | 0.27 μg.g^-1^  95% CI [0.23, 0.34] | r = 0.84 (large)  95% CI [0.74, 0.85] |
| *Acanthurus bahianus* | Muscle | Wilcoxon | W = 100 | Ø | < .001 | 20 | 0.63 μg.g^-1^  95% CI [0.39, 0.80] | r = 0.84 (large)  95% CI [0.74, 0.85] |
| *Diadema antillarum* | Muscle | Student (Welch correction) | t = 6.99 | 9.00 | < .001 | 20 | 1.57 μg.g^-1^  95% CI [1.06, 2.08] | d = 3.12 (large)  95% CI [1.73, 4.53] |
| *Diadema antillarum* | Gonad | Wilcoxon | W = 100 | Ø | < .001 | 20 | 4.24 μg.g^-1^  95% CI [2.64, 5.79] | r = 0.84 (large)  95% CI [0.74, 0.85] |
| *Abudefduf saxatilis* | Liver | Wilcoxon | W = 90 | Ø | < .001 | 19 | 45.1 μg.g^-1^  95% CI [12.8, 85.7] | r = 0.78 (large)  95% CI [0.61, 0.85] |
| *Acanthurus bahianus* | Liver | Wilcoxon | W = 88 | Ø | < .001 | 19 | 70.9 μg.g^-1^  95% CI [52.6, 118] | r = 0.82 (large)  95% CI [0.67, 0.85] |

**Table S2** – Statistical comparison of mercury concentrations in Bouillante and in the Control Site for each type of samples.

**Table S3** – Maximum Safe Weekly Consumption (MSWC) with 95% CI under both mercury form scenario, calculated after the mean concentrations and their robust 95% CI (Table S1). As a guideline for interpretation, gross approximations of the number of adults containing 100g of each tissue are given.

| Species | Tissue | Number of individuals per 100g of tissue | Zone | Hg form | MSWC | Unit |
| --- | --- | --- | --- | --- | --- | --- |
| *Diadema antillarum* | Gonad | ~12.5 adults  Note: Based the weigth of gonads of *Echinometra lucunter* (another guadeloupean sea urchin), which is similarly sized (8g; Ebert et al., 2008) | Control | Inorganic | 2430.4  95% CI [1850.8, 3353.1] | g.week^-1^ |
|  |  |  |  | MeHg | 972.2  95% CI [740.3, 1341.2] |  |
|  |  |  | Bouillante | Inorganic | 167.2  95% CI [77.9, 282.9] |  |
|  |  |  |  | MeHg | 66.9  95% CI [31.2, 113.2] |  |
| *Acanthurus bahianus* | Muscle | ~2 adults  Note: Considering that muscle is 50% of the weight in fishes (Gremyachikh et al., 2018). Calculated for the mean weight of adults: (100g; Robertson et al., 2005; Macieira & Joyeux, 2008). | Control | Inorganic | 83.6  95% CI [45.3, 108.6] | kg.week^-1^ |
|  |  |  |  | MeHg | 33.4  95% CI [18.1, 43.5] |  |
|  |  |  | Bouillante | Inorganic | 1.7  95% CI [1.2, 2.6] |  |
|  |  |  |  | MeHg | 0.7  95% CI [0.5, 1.0] |  |
| *Spondylus tenuis* | Muscle | ~6 adults  Note: Based on *S. calcifer* adductor muscle weight (32.5g; Cudney-Bueno & Rowell, 2008), which is around twice bigger than *S. tenuis* (Rocha et al., 2015). | Control | Inorganic | 30.2  95% CI [23.1, 34.0] | kg.week^-1^ |
|  |  |  |  | MeHg | 12.1  95% CI [9.2, 13.6] |  |
|  |  |  | Bouillante | Inorganic | 3.1  95% CI [1.0, 7.8] |  |
|  |  |  |  | MeHg | 1.2  95% CI [0.4, 3.1] |  |
